# Rewiring of aminoacyl-tRNA synthetase localization and interactions in plants with extensive mitochondrial tRNA gene loss

**DOI:** 10.1101/2022.01.27.478071

**Authors:** Jessica M. Warren, Amanda K. Broz, Ana Martinez-Hottovy, Christian Elowsky, Alan C. Christensen, Daniel B. Sloan

## Abstract

The number of tRNAs encoded in plant mitochondrial genomes varies considerably. Ongoing loss of bacterial-like mitochondrial tRNA genes in many lineages necessitates the import of nuclear-encoded eukaryotic counterparts that share little sequence similarity. Because tRNAs are involved in highly specific molecular interactions, this replacement process raises questions about the identity and trafficking of enzymes necessary for the maturation and function of newly imported tRNAs. In particular, the aminoacyl-tRNA synthetases (aaRSs) that charge tRNAs are usually divided into distinct classes that specialize on either organellar (mitochondrial and plastid) or cytosolic tRNAs. Here, we investigate the evolution of aaRS subcellular localization in a plant lineage (*Sileneae*) that has experienced extensive and rapid mitochondrial tRNA loss. By analyzing full-length mRNA transcripts (PacBio Iso-Seq), we found the predicted retargeting of many ancestrally cytosolic aaRSs to the mitochondrion and confirmed these results with colocalization microscopy assays. However, we also found cases where aaRS localization does not appear to change despite functional tRNA replacement, suggesting evolution of novel interactions and charging relationships. Therefore, the history of repeated tRNA replacement in *Sileneae* mitochondria reveals that differing constraints on tRNA/aaRS interactions may determine which of these alternative coevolutionary paths is used to maintain organellar translation in plant cells.

## Introduction

Translation in the plant cell is a tripartite system. The presence of a nuclear and two organellar (plastid and mitochondrial) genomes results in protein synthesis occurring in three separate compartments. Although the bacterial progenitors of plastids and mitochondria harbored all genetic components required for translation, their genomes have since been extensively reduced, and numerous proteins involved in organellar translation are now encoded in the nucleus and imported into the organelles (Huang et al., 2003; Timmis et al., 2004; Giannakis et al., 2022). Transfer RNAs (tRNAs) are some of the last remaining translational components encoded in organellar genomes. Most bilaterian animals contain a minimally sufficient set of mitochondrial tRNA genes (Boore, 1999), but the number of tRNAs encoded in plant mitogenomes can vary dramatically. Some angiosperm mitogenomes even exhibit rapid and ongoing mitochondrial tRNA gene loss within single genera (Sloan et al., 2012b; Petersen et al., 2015). Loss of these bacterial-like tRNAs inherited from the endosymbiotic ancestor of mitochondria necessitates the import of nuclear-encoded (eukaryotic) tRNAs to maintain mitochondrial protein synthesis (Salinas-Giegé et al., 2015). The import of nuclear tRNAs into plant mitochondria has been recognized for decades (Small et al., 1992; Delage et al., 2003), but there are longstanding questions about how tRNA import evolves. In particular, which enzymes are responsible for the maturation and function of these imported tRNAs, and how has their subcellular trafficking evolved in association with changes in tRNA import? The enzymes that recognize tRNAs and charge them with the correct amino acid are known as aminoacyl-tRNA synthetases (aaRSs) and are usually divided into two distinct classes that specialize on either organellar or cytosolic tRNAs. In most eukaryotes, including vascular plants, all aaRS genes are encoded by the nuclear genome (Duchêne et al., 2009). Therefore, aaRSs that function in organellar protein synthesis must be translated by cytosolic ribosomes, targeted to the correct organelle, and translocated across multiple membranes (Duchêne et al., 2009; Ghifari et al., 2018). These organellar aaRSs largely originate from intracellular gene transfers (plastid and mitochondrial transfers to the nuclear genome) or horizontal gene transfers from other bacterial sources, making them highly divergent from their cytosolic counterparts (Doolittle and Handy, 1998; Duchêne et al., 2005; Brandao and Silva-Filho, 2011; Rubio Gomez and Ibba, 2020).

The import of nuclear-encoded aaRSs into plant organelles is primarily achieved through amino acid sequences at their N-termini (transit peptides) that are recognized by translocase proteins on outer organelle membranes (Berglund et al., 2009; Ge et al., 2014; Ghifari et al., 2018). These transit peptides can vary considerably in length from fewer than 20 amino acids to over 100 (averaging around 42-50 residues) and are cleaved after translocation across the organellar membranes (Huang et al., 2009; Ge et al., 2014; Murcha et al., 2014). Mitochondrial transit peptides often form amphipathic alpha helices with alternating hydrophobic and positively charged amino acids (Huang et al., 2009; Schmidt et al., 2010). Plant mitochondrial transit peptides are also particularly rich in Ser residues, and many have a loosely conserved motif containing an Arg residue near the peptide cleavage site (Huang et al., 2009; Ge et al., 2014). Despite these general structural features, there is very little primary amino acid sequence conservation in transit peptides (Lee et al., 2008; Kunze and Berger, 2015), and these domains are considered some of the fastest evolving (non-neutral) sites (Williams et al., 2000; Christian et al., 2020).

Somewhat surprisingly, analyses of aaRS genes in *Arabidopsis* thaliana did not find the expected 20 aaRS (one aaRS for each proteinogenic amino acid) genes for each subcellular compartment (cytosol, mitochondria, and plastids) (Small et al., 1999; Duchêne et al., 2005). Instead, most organellar aaRSs function in both mitochondria and plastids — reducing the number of aaRSs in *A. thaliana* to only 45 (Duchêne et al., 2005). These dual-targeted aaRSs must then interact with both mitochondrial tRNAs (mt-tRNAs) and plastid tRNAs to enable translation in these bacterial-like systems.

Dual-targeted aaRSs that function in both mitochondria and plastids contain an ambiguous N-terminal transit peptide that is recognized by both organelle outer membranes (Peeters and Small, 2001; Duchêne et al., 2005). While plastid-specific transit peptide sequences generally lack the helical structure found on mitochondrial transit peptides, both organelle transit peptides have very similar amino acid compositions with many hydrophobic and positively charged residues (Bruce, 2001; Ge et al., 2014; Christian et al., 2020). Not surprisingly, dual-targeted transit peptides often exhibit intermediate properties between plastid- and mitochondrial-specific transit peptides (Pujol et al., 2007; Berglund et al., 2009).

Although most of the aaRSs imported into plant organelles are dual-targeted and bacterial-like, there are exceptions. In *A. thaliana*, five eukaryotic-like (cytosolic) aaRSs are dual-localized to mitochondria and the cytosol (Mireau et al., 1996; Duchêne et al., 2005). The import of these eukaryotic-like aaRSs demonstrates the complex nature of mitochondrial tRNA metabolism in plants, where the import of some nuclear-encoded tRNAs is also necessary because the mitogenome contains an incomplete set of tRNAs (Michaud et al., 2011). The five aaRS enzymes shared between the cytosol and mitochondria in *A. thaliana* correspond to tRNAs that are also imported from the cytosol – thereby maintaining phylogenetic congruence between the imported tRNA and interacting enzyme (Duchêne et al., 2005). This coevolutionary pairing of tRNAs and aaRSs may be necessary due to the highly discriminating nature of aaRSs (Rubio Gomez and Ibba, 2020). The attachment of the correct amino acids to corresponding tRNAs is essential for the faithful decoding of the genome and is achieved through a highly accurate process whereby aaRS enzymes use certain nucleotide positions (identity elements) on the tRNA for substrate recognition (Giege et al., 1998). As eukaryotic tRNAs have little sequence similarity with the bacterial-like mitochondrial and plastid tRNAs, they would be expected to make poor substrates for bacterial-like aaRSs (Salinas-Giegé et al., 2015).

However, there are cases of an aaRS and tRNA that functionally interact despite originating from different domains of life (Duchêne et al., 2005; Warren and Sloan, 2020). For example, a eukaryotic ProRS appears to have replaced all bacterial counterparts in the genome of *A. thaliana*, despite the species retaining a bacterial-like mitochondrial tRNA-Pro. Therefore, mitochondrial tRNA-Pro must then be charged by a eukaryotic enzyme. However, two eukaryotic ProRSs exist in the *A. thaliana* genome, and only one of those genes contains an organellar transit peptide – suggesting that some enzymatic differentiation may be necessary for bacterial-like tRNA recognition (Duchêne et al., 2005).

Despite a few aaRS/tRNA phylogenetic incongruencies, there exists a general rule of tRNAs encoded in the mitogenome being charged by enzymes that are bacterial in nature. Questions then arise as to the trafficking of aaRSs in plants that have undergone recent and extensive mt-tRNA loss. For example, mitogenomes from close relatives within the angiosperm tribe *Sileneae* exhibit a wide range of mt-tRNA gene content (Fig. 1) (Sloan et al., 2010; Sloan et al., 2012b), and recent analysis indicates that these mt-tRNAs in this lineage are being functionally replaced by import of nuclear-encoded counterparts (Warren et al., 2021).

**Figure 1.**
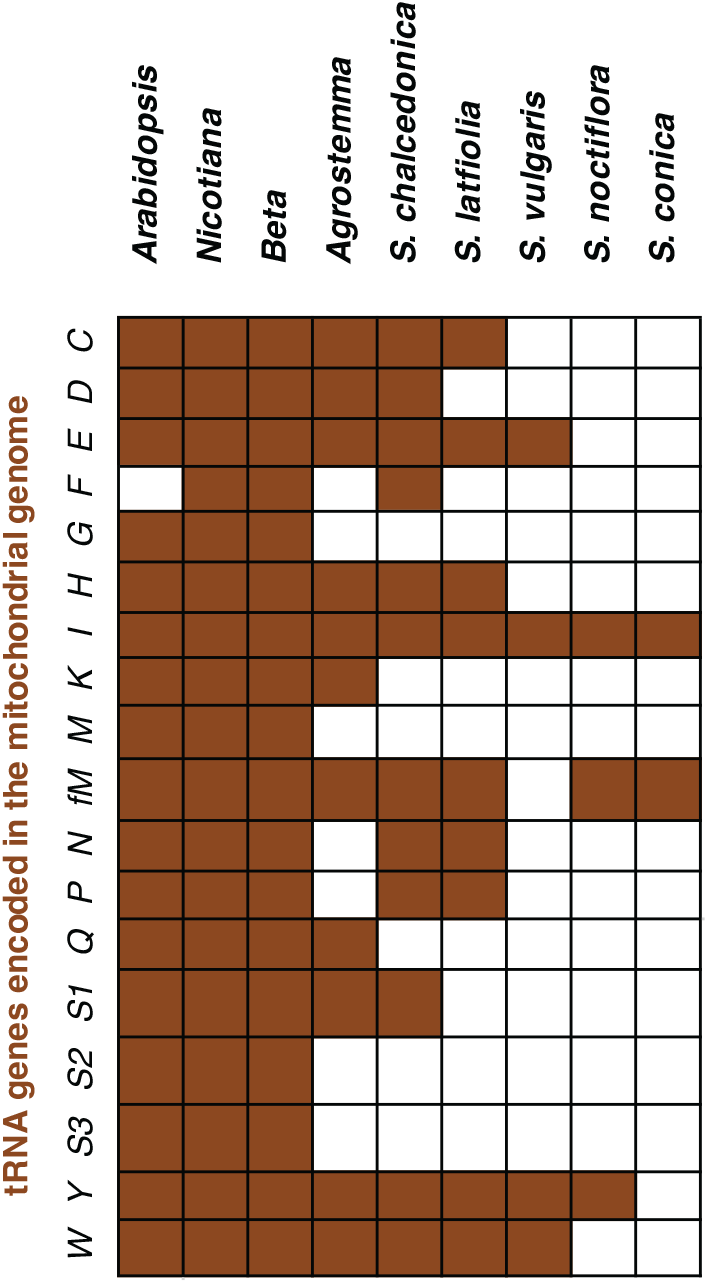
The loss of mt-tRNA genes in the mitogenomes of multiple *Sileneae* species. Singe-letter codes are used for amino acid abbreviations. Filled squares indicate the presence of the corresponding tRNA gene in the mitogenome.

The almost complete loss and replacement of native mt-tRNAs with nuclear-encoded tRNAs in *Sileneae* species raises multiple alternative scenarios as to the identity of the aaRSs that aminoacylate these newly imported tRNAs (Fig. 2). It is possible that the ancestrally cytosolic aaRSs evolved de novo targeting to the mitochondria and act on the newly imported tRNAs – effectively replacing both partners in the mitochondrial tRNA/aaRS system with cytosolic counterparts (Fig. 2A). Alternatively, the ancestral organellar aaRSs could retain mitochondrial localization and now recognize novel substrates (cytosolic tRNAs), either through adaptation or preexisting enzymatic promiscuity (Fig. 2B). In this study, we test for these alternative hypotheses in the angiosperm clade *Sileneae* to gain insight into the cellular and molecular mechanisms that facilitate the loss and functional replacement of mitochondrial tRNA genes in plants. By using full-length mRNA sequencing and fluorescent co-localization microscopy, we show that both evolutionary scenarios are likely at play with roughly equal frequency in systems rapidly losing mitochondrial tRNAs. We also found evidence that perturbation of an aaRS/tRNA interaction in mitochondria may have pleiotropic effects on plastid aaRS evolution. And finally, we offer a possible explanation as to why the retargeting of an ancestrally cytosolic aaRS may be necessary in some, but not all, cases of tRNA replacement by exploring known identity elements in these aaRS/tRNA interactions.

**Figure 2.**
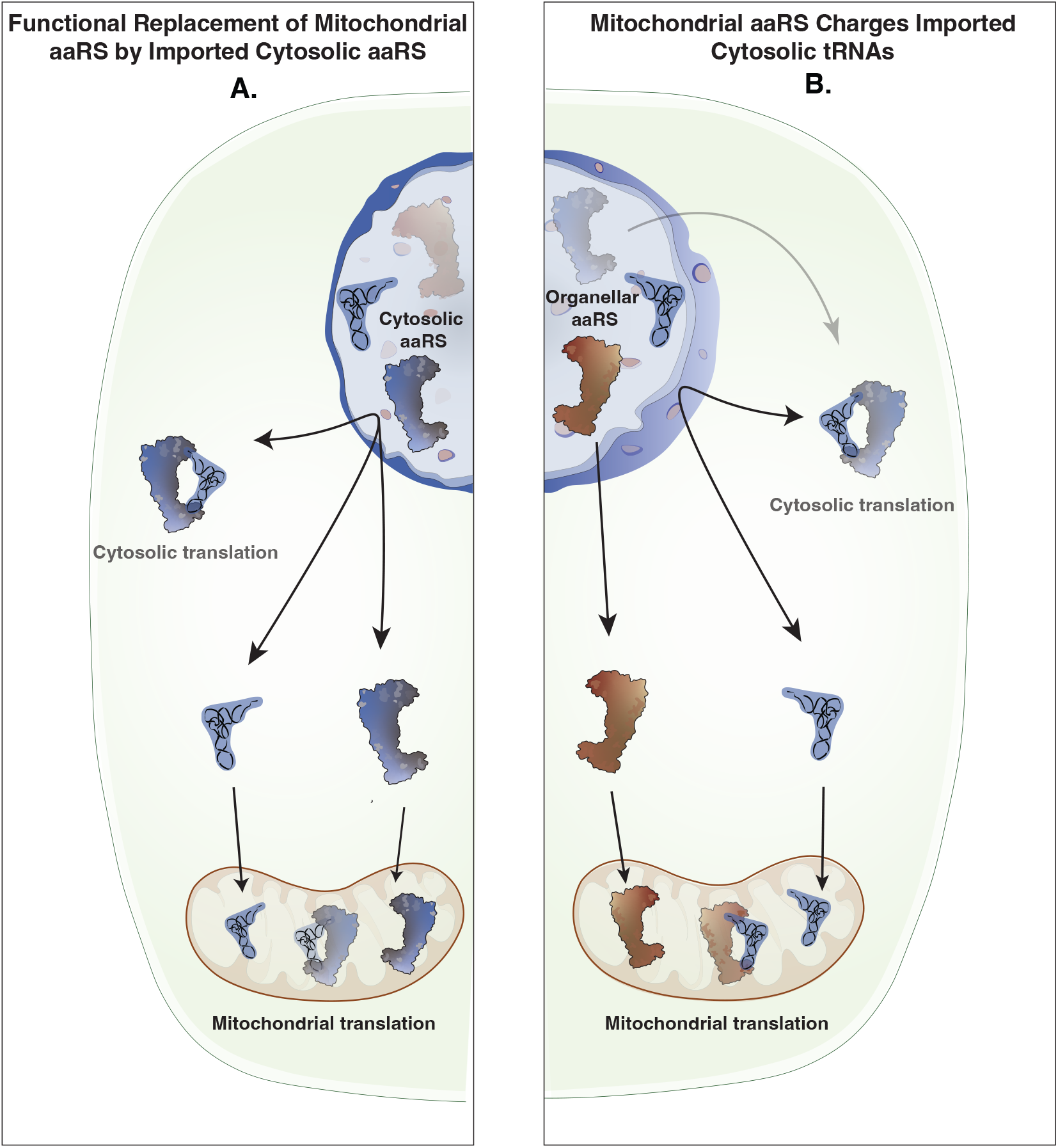
Alternative scenarios for the aminoacylation of nuclear tRNAs imported into *Sileneae* mitochondria. Panel (A) shows the import of an aaRS that was ancestrally only found in the cytosol but is now trafficked to mitochondria to charge the corresponding nuclear encoded, imported tRNAs. Panel (B) illustrates the retention of the ancestral trafficking of the organellar enzyme which now must aminoacylate the newly imported tRNA – creating a phylogenetic mismatch between aaRS and tRNA.

## Results and Discussion

### Identification and characterization of Sileneae aaRS gene content

Putative transit peptides can be identified with prediction programs that search for characteristic secondary structure, amino acid composition, and peptide cleavage-site motifs (Small et al., 2004; Sperschneider et al., 2017; Almagro Armenteros et al., 2019). To test for the gain of organellar transit peptides on ancestrally cytosolic aaRSs in *Sileneae* species, we sequenced full-length mRNA transcripts from five species (*Agrostemma githago, Silene conica, S. latifolia, S. noctiflora, and S. vulgaris*), using PacBio Iso-Seq technology (Zhao et al., 2019). Full-length mRNA sequences are useful when inferring which specific gene copies have N-terminal extensions because plants often have multicopy genes with high sequence similarity. Previously generated genome assemblies from the same species (Krasovec et al., 2018; Warren et al., 2021; Williams et al., 2021) were also searched for genes and putative transit peptides potentially missed by Iso-Seq analysis due to lower expression levels.

This analysis identified transcripts from each *Sileneae* species corresponding to known *A. thaliana* organellar and cytosolic aaRSs for each amino acid (Supp. Table 1). As expected, *Sileneae* aaRSs that were homologous to organelle-targeted aaRSs in *A. thaliana* had very high predicted probabilities of being localized to mitochondria, plastids, or both (Supp. Figs. 1-20). However, multiple cytosolic aaRS genes that lack transit peptides in *A. thaliana* had N-terminal extensions in one or more *Sileneae* species.

### Mt-tRNA loss in Sileneae is associated with frequent acquisition of putative aaRS transit peptides

In *Sileneae*, mt-tRNA genes decoding 13 amino acids have been lost in one or more species compared to *A. thaliana*, and a 14th (mt-tRNA-Phe) was lost independently in *A. thaliana* and some *Sileneae* species (Fig. 1). These 14 losses raise the question as to which aaRSs are charging the newly imported cytosolic tRNAs that have functionally replaced these mt-tRNAs. In seven of these cases, an N-terminal extension predicted to serve as a mitochondrial transit peptide was found on a cytosolic aaRS in multiple *Sileneae* species: GlnRS (Fig. 3A), GlyRS (Supp. Fig. 8), LysRS (Fig. 4A), TyrRS (Fig. 5A), MetRS, ProRS, TrpRS (Fig. 6A-C). In these cases, the corresponding *A. thaliana* enzyme is not mitochondrial-targeted, implying evolutionary gains of transit peptides and targeting in *Sileneae*. These examples of aaRS retargeting indicate that ancestral pairings between cytosolic aaRSs and cytosolic tRNAs are maintained and have expanded their function to include mitochondrial translation. Duplication and gain of function is a common theme in protein evolution (Lynch, 2007) and likely played a role in the mitochondrial targeting of ancestrally cytosolic aaRSs in *Sileneae*. We found that many aaRS genes existed as multicopy gene families, and there were multiple cases where an N-terminal extension was only present in one of the gene copies within an aaRS family: GlnRS (Fig. 3A), TyrRS (Fig. 5A), ProRS (Fig. 6A) and MetRS (Fig. 6C). In these cases, it appears that mitochondrial localization happened following a gene duplication event. The age of these duplications varied considerably, as the two groups of cytosolic MetRS enzymes predate the divergence of *A. thaliana* and *Sileneae* (see Supp. Fig. 13 for MetRS1), whereas the duplication of GlnRS, TyrRS and ProRS was specific to the lineage leading to *Sileneae* (Figs. 4A, 6A, 7A). TrpRS was the only one of the cytosolic aaRS enzymes predicted to gain a mitochondrial transit peptide that was clearly present as a single copy in Silene (Fig. 7B).

**Figure 3.**
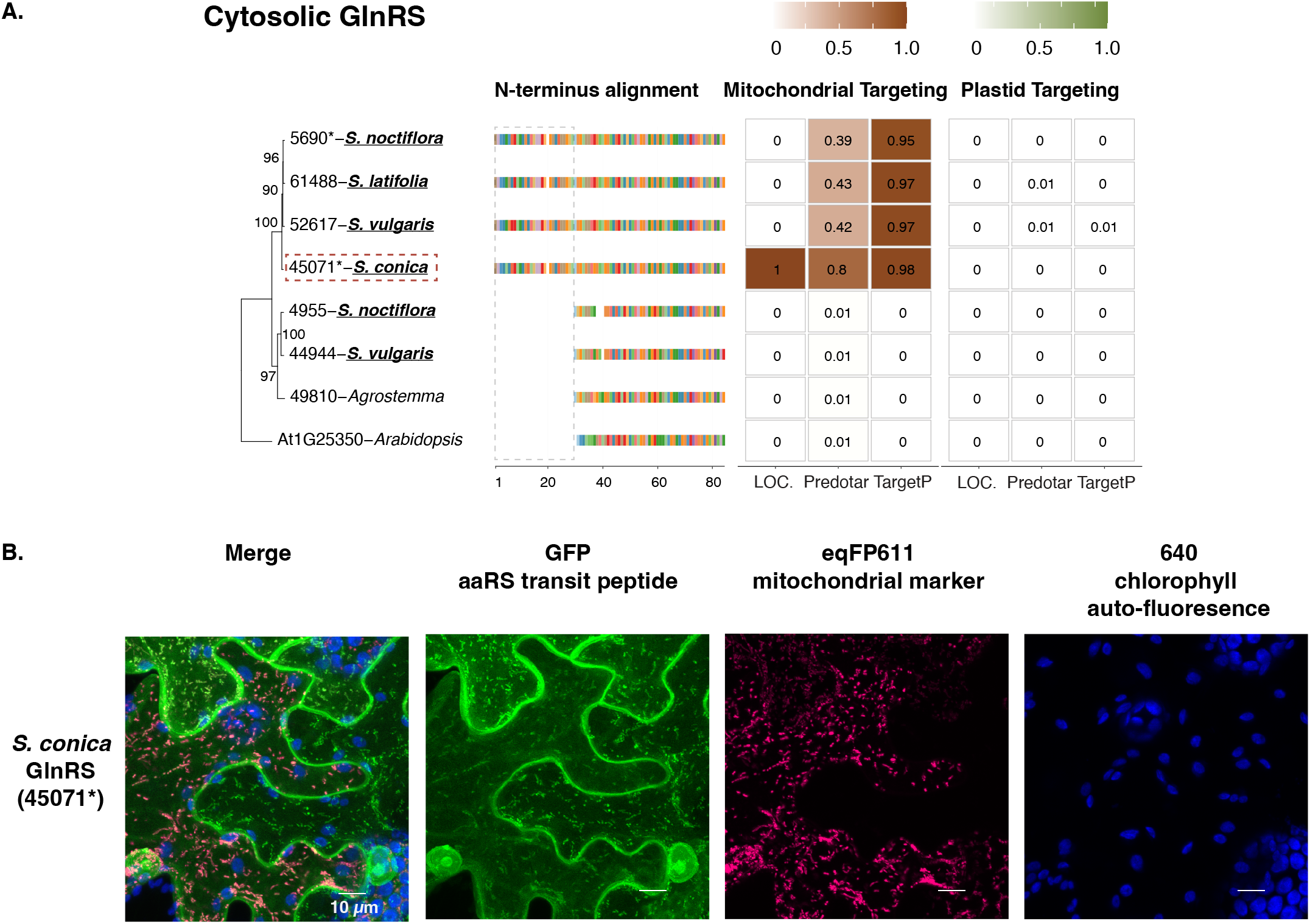
Cytosolic GlnRS enzymes in *Sileneae* species have N-terminal extensions that are predicted to function as mitochondrial transit peptides and can target to mitochondria in *N. benthamiana*. (A) A maximum likelihood tree of the complete enzyme sequences from *A. thaliana* and *Sileneae* species is shown with bootstrap values indicated. To the right of the tree is an alignment of just the first 84 amino acids of the aaRS proteins. The color key for the amino acids can be found in Supp. Fig. 21. The gray dashed box highlights the difference in N-terminal length between the clade of enzymes found in *Silene* and the rest of the sequences. Right of the alignment are the targeting probabilities generated from the respective software program for each GlnRS gene. Genes marked with an * indicate that the gene was detected in Iso-Seq data but without the 5’ extensions present in the nuclear assembly. Species underlined in bold font have lost the corresponding mt-tRNA gene from the mitochondrial genome and must now import a nuclear tRNA counterpart. The predicted transit peptide from the gene highlighted with a red dashed box was fused to GFP for the analysis in panel B. (B) Transient expression of the predicted transit peptide from gene 45071 in *N. benthamiana* epithelial cells. The amino acid sequence plus 10 upstream amino acids of the protein body were fused to GFP and co-transfected with an eqFP611-tagged transit peptide from a known mitochondrially localized protein (isovaleryl-CoA dehydrogenase). All scale bars represent 10 *µ*M.

**Figure 4.**
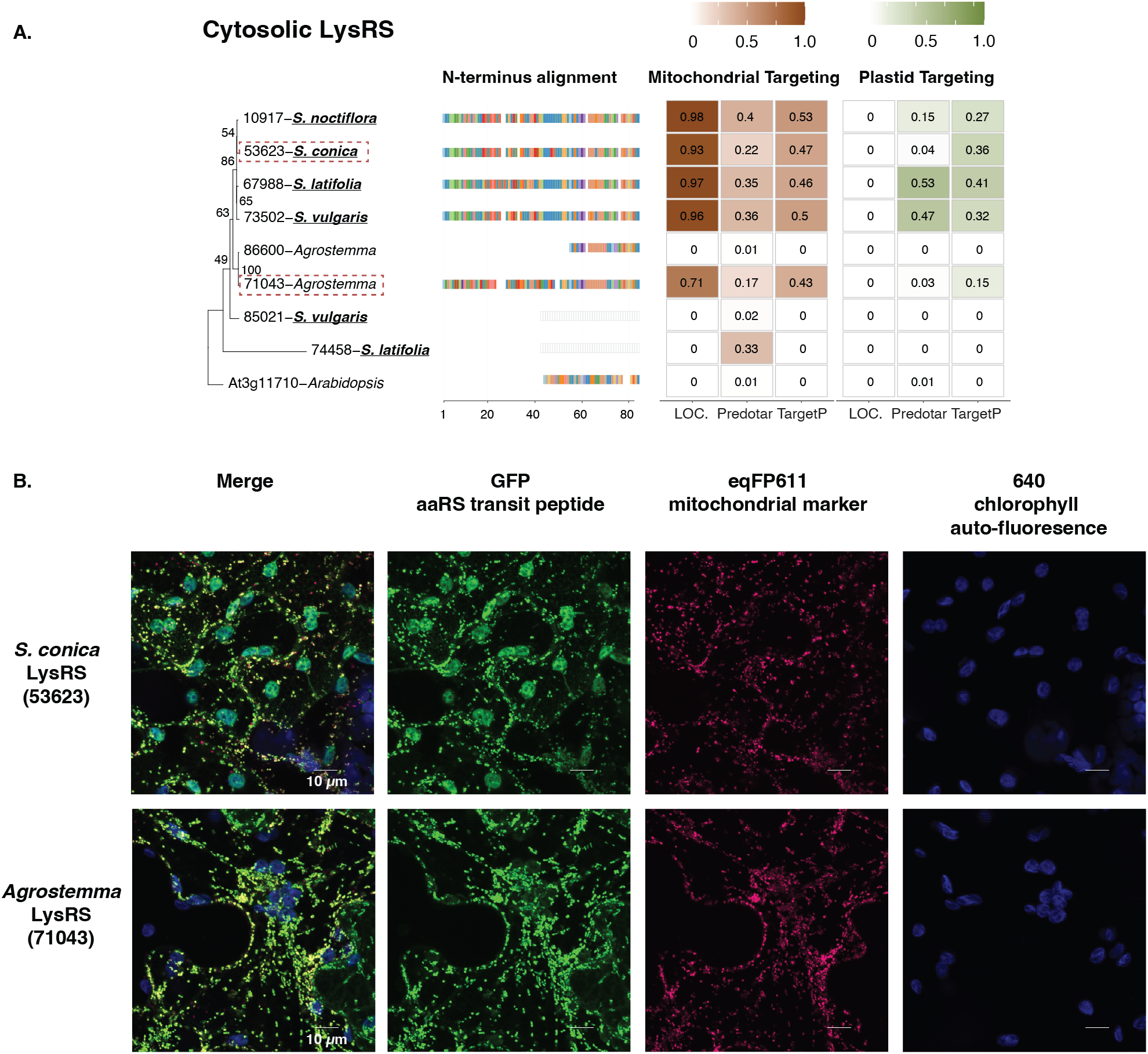

**Cytosolic LysRS enzymes in *Sileneae* species have N-terminal extensions that are predicted to function as mitochondrial transit peptides and can target mitochondria in** *N. benthamiana*.

(A) A maximum likelihood tree of the complete enzyme sequences from *A. thaliana* and *Sileneae* species is shown with bootstrap values indicated. To the right of the tree is an alignment of just the first 84 amino acids of the aaRS proteins. The color key for the amino acids can be found in Supp. Fig. 21. Right of the alignment are the targeting probabilities generated from the respective software program for each LysRS gene. Species underlined in bold font have lost the corresponding mt-tRNA gene from the mitochondrial genome and must now import a nuclear tRNA counterpart. The predicted transit peptide from the gene highlighted with a red dashed box was fused to GFP for the analysis in panel B. (B) Transient expression of the predicted transit peptide from gene 45071 in *N. benthamiana* epithelial cells. The amino acid sequence plus 10 upstream amino acids of the protein body were fused to GFP and co-transfected with an eqFP611-tagged transit peptide from a known mitochondrially localized protein (isovaleryl-CoA dehydrogenase). All scale bars represent 10 *µ*M.

**Figure 5.**
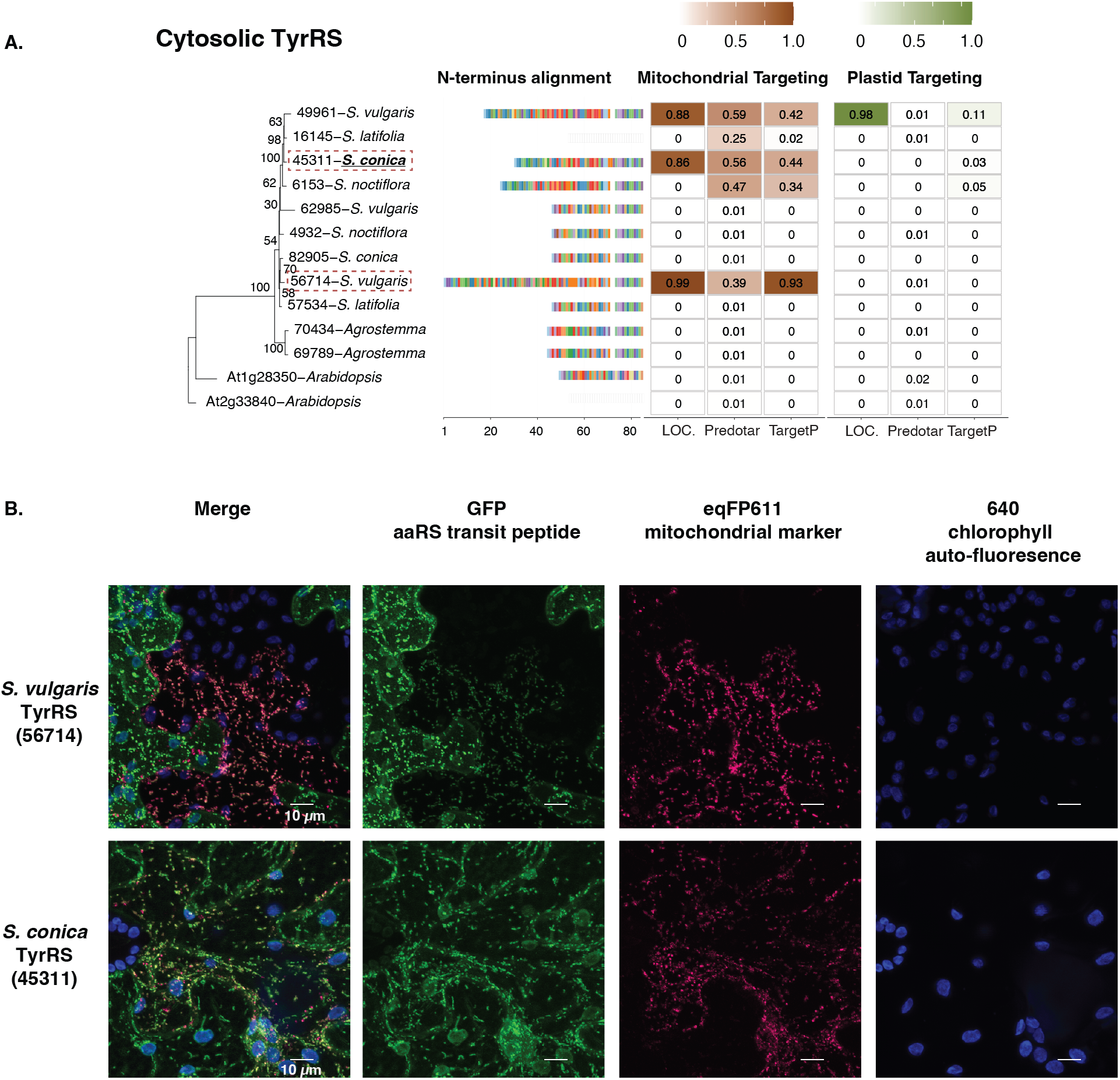
Cytosolic TyrRS enzymes in *Sileneae* species have N-terminal extensions that are predicted to function as mitochondrial transit peptides and can target mitochondria in *N. benthamiana*. (A) A maximum likelihood tree of the complete enzyme sequences from *A. thaliana* and *Sileneae* species is shown with bootstrap values indicated. To the right of the tree is an alignment of just the first 84 amino acids of the aaRS proteins. The color key for the amino acids can be found in Supp. Fig. 21. Right of the alignment are the targeting probabilities generated from the respective software program for each TrpRS gene. Species underlined in bold font have lost the corresponding mt-tRNA gene from the mitochondrial genome and must now import a nuclear tRNA counterpart. The predicted transit peptide from the gene highlighted with a red dashed box was fused to GFP for the analysis in panel B. (B) Transient expression of the predicted transit peptide from gene 45071 in *N. benthamiana* epithelial cells. The amino acid sequence plus 10 upstream amino acids of the protein body were fused to GFP and co-transfected with an eqFP611-tagged transit peptide from a known mitochondrially localized protein (isovaleryl-CoA dehydrogenase). All scale bars represent 10 *µ*M.

**Figure 6.**
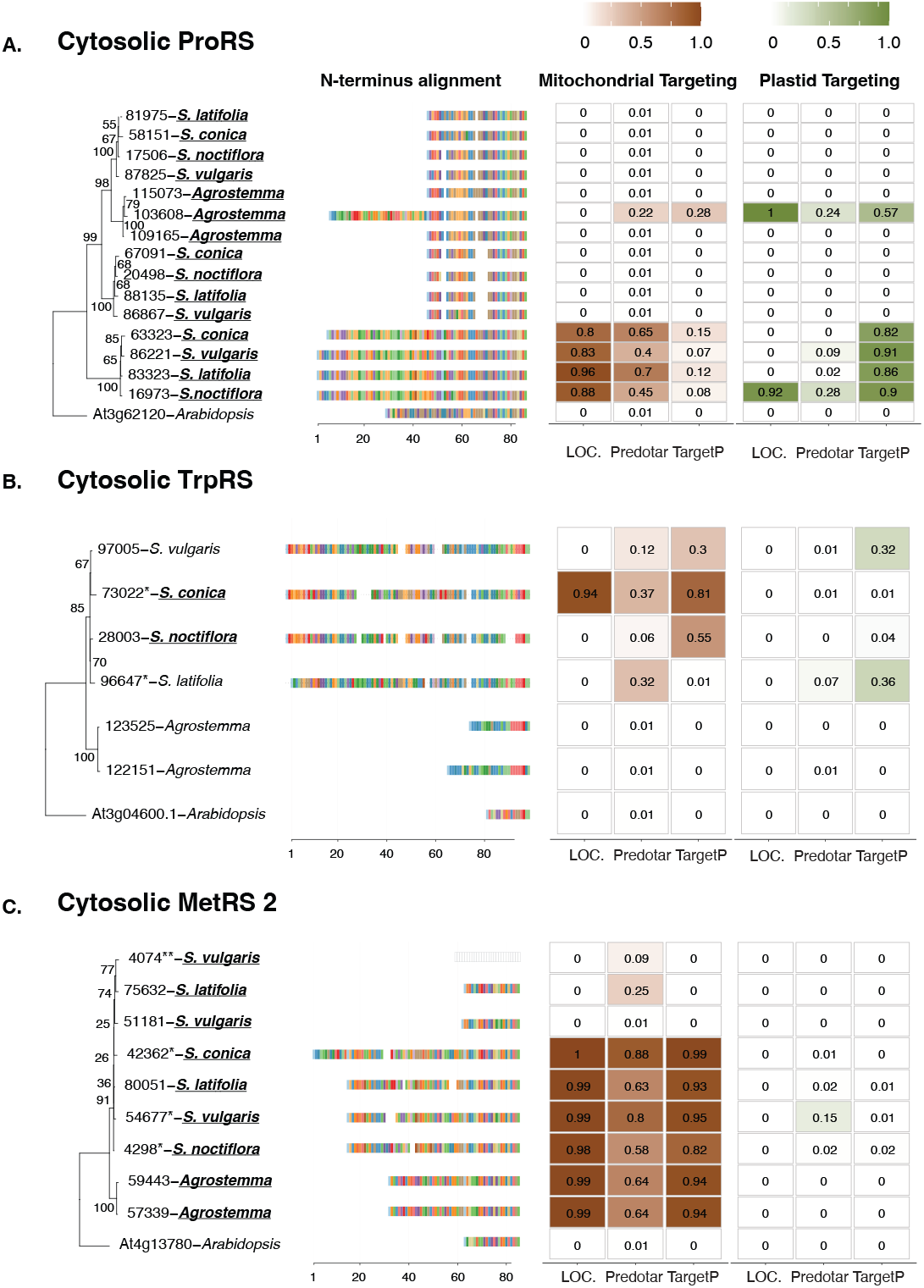
Predicted targeting and N-terminal alignments of cytosolic ProRS, TrpRS, and MetRS2 in multiple *Sileneae* species and *A. thaliana*. Each panel represents a maximum likelihood tree of the complete aaRS sequences from *A. thaliana* and *Sileneae* species, with bootstrap values indicated on branches. To the right of the tree is an alignment of just the first 84 amino acids of the aaRS proteins. The color key for the amino acids can be found in Supp. Fig. 21. Right of the alignment are the targeting probabilities generated from the respective software program for each aaRS. Species underlined in bold font have lost the corresponding mt-tRNA gene from the mitochondrial genome and must now import a nuclear tRNA counterpart.

**Figure 7.**
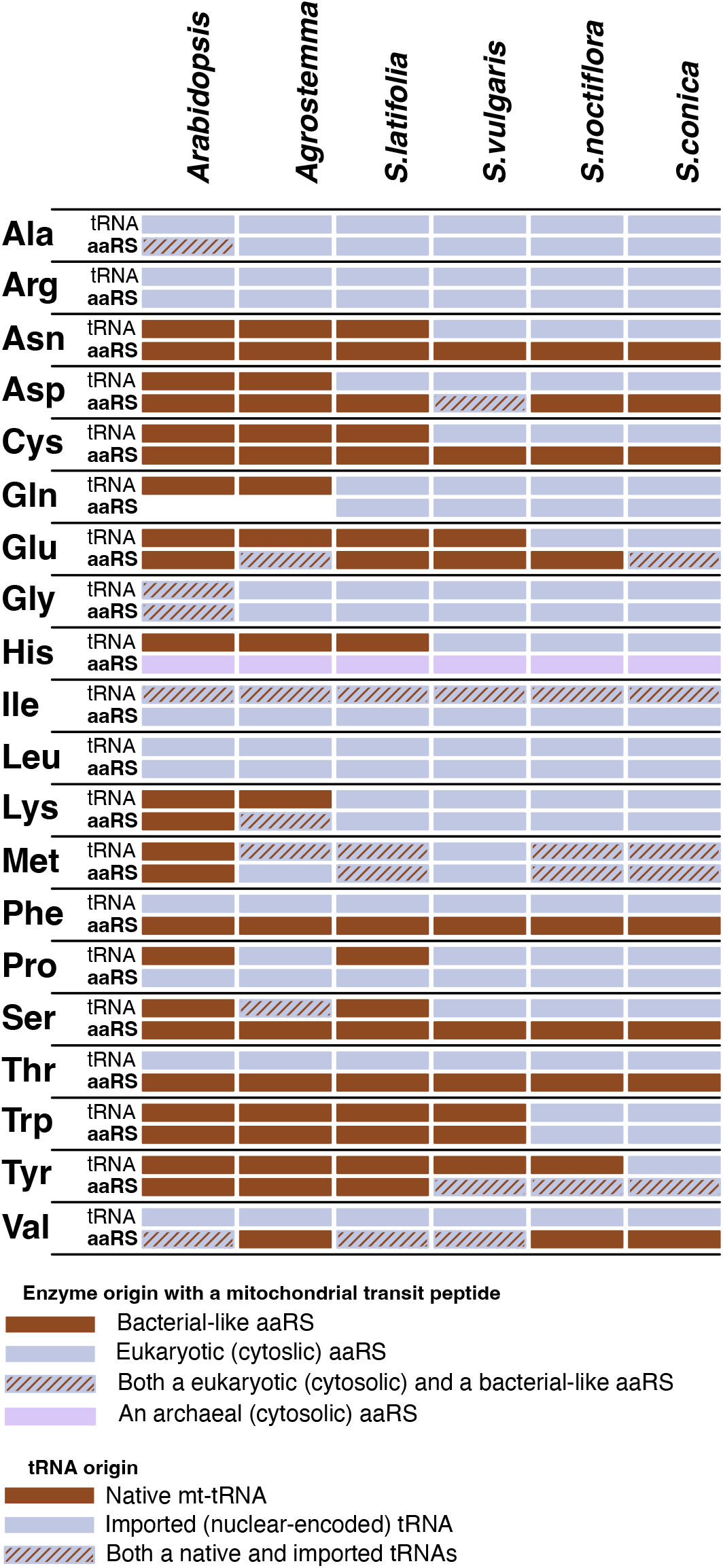
The origin of mitochondrial tRNAs and the cognate mitochondrially localized aaRSs in *A. thaliana* and multiple *Sileneae* species. The mitochondrial localization of aaRSs has been experimentally demonstrated in *A. thaliana* (see Supp. Table 1, (Duchêne et al., 2005)) and has been assigned for Silenae species based on localization prediction software. Mt-tRNAs and aaRSs have complex gene histories. Thus, tRNAs and aaRSs classified as “bacterial-like” include those derived from the mitochondria (alphaproteobacterial-like), the plastids (cyanobacterial-like), or other bacterial origins. All bacterial-like tRNAs are still encoded in the mitogenomes, whereas all eukaryotic mitochondrial tRNAs are imported from the cytosol. Solid colors represent the presence of only a single phylogenetic class of a tRNA or aaRS, while striped boxes indicate the presence of multiple aaRSs/tRNAs with differing phylogenetic origins. The cytosolic TyrRS in angiosperms is archaeal in origin, and it appears to have been retargeted to the mitochondria in some *Silene* species. For this figure, an aaRS was classified as being predicted to be mitochondrially targeted if the enzyme had 50 or more percentage points of targeting likelihood to the organelle (cumulatively between the three targeting prediction software programs). See Supp. Figs 1-20 for more detailed targeting data for each gene family. An organellar GlnRS does not exist in most plant mitochondria (including *Arabidopsis*) as tRNA-Gln is aminoacylated by an indirect transamidation pathway with a nondiscriminating GluRS (Salinas-Giegé et al., 2015). This ancestral state appears to be retained in *Agrostemma*. All ProRS enzymes have eukaryotic origins, but organelle and cytosolic genes are distinct.

There were also cases where mitochondrial localization was associated with alternative transcription start sites that resulted in the expression of two isoforms – one with and one without an N-terminal extension predicted to be a transit peptide. Presumably, the isoforms without the extensions have retained their ancestral function in the cytosol. For MetRS, GlnRS, LysRS, and TrpRS expression, the isoform lacking an N-terminal extension (but otherwise identical or nearly identical to the extension-containing transcripts) exhibited much higher expression levels (inferred from Iso-Seq read counts) than the isoform with a predicted transit peptide.

### The N-terminal extensions found on Sileneae aaRS enzymes can confer mitochondrial targeting in Nicotiana benthamiana

To test whether the N-terminal extensions found on aaRS transcripts could function as mitochondrial transit peptides, the entire transit peptide region predicted by TargetP v.2.0 (Almagro Armenteros et al., 2019) plus 10 amino acids of the protein coding body was fused to the 5′-end of green fluorescent protein (GFP) and co-infiltrated with a mitochondrial-targeted red fluorescent protein eqFP611 into *Nicotiana benthamiana* epidermal leaf cells.

GFP constructs with predicted transit peptides were made for eight genes in total, one for GlnRS (Fig. 3B), two for LysRS (Fig. 4B), two for TyrRS (Fig. 5B), and three for PheRS (Fig. 8 C-D). All peptides tested exhibited a strong mitochondrial GFP/eqFP611 colocalization signal confirming that these amino acid sequences could be used to target proteins to plant mitochondria.

Somewhat surprisingly, the N-terminal extensions accumulation in chloroplasts to varying degrees (Fig. 4B and Fig. 5B). Transient expression of the construct containing the N-terminal extension of GlnRS also resulted in membrane and nuclearaccumulationofGFP(inadditiontoastrongmitochondrial localization signal) but did not localize to chloroplasts (Fig. 3B).

**Figure 8.**
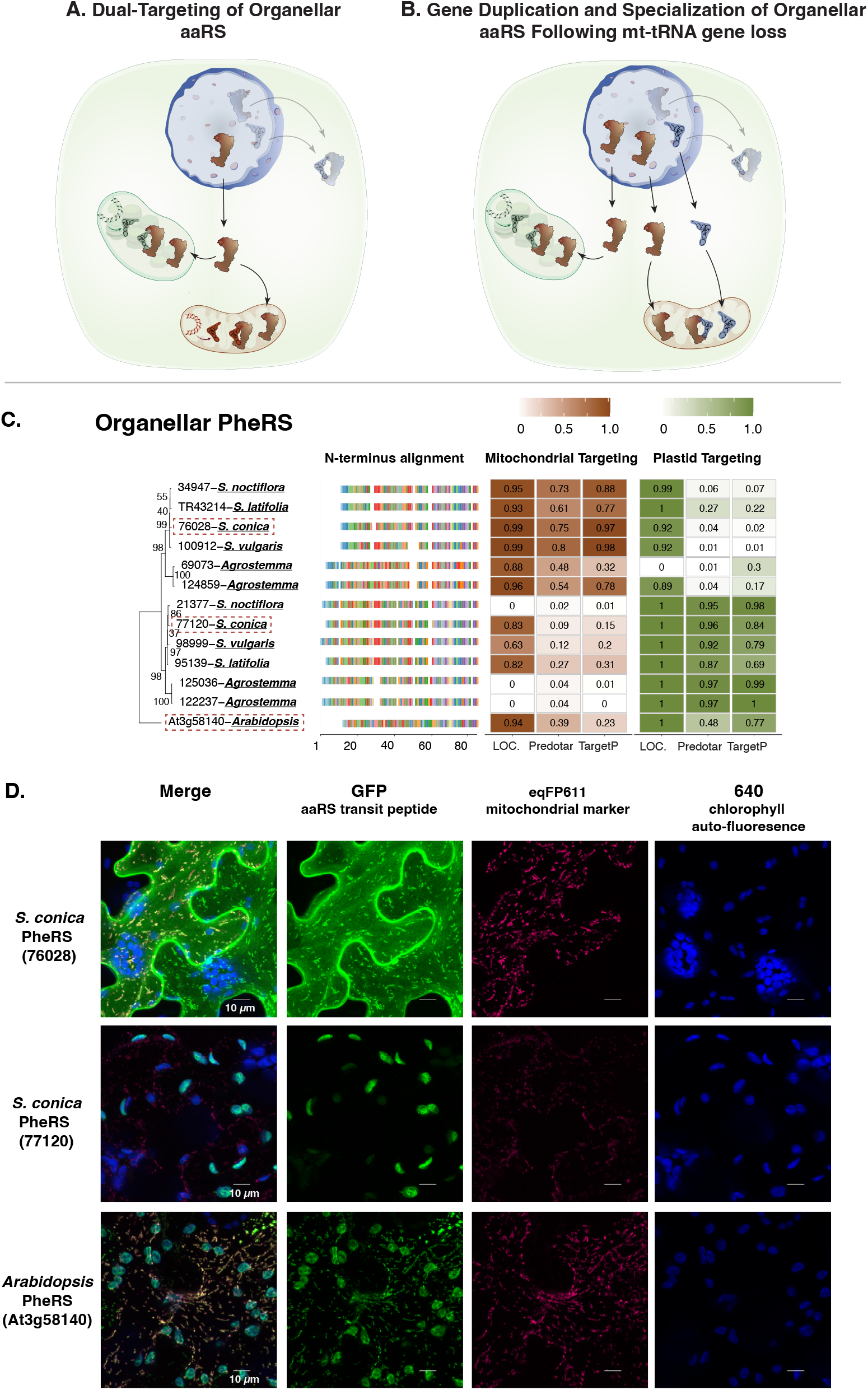
Loss of the mitochondrial tRNA-Phe precipitates duplication and subfunctionalization of the organellar PheRS enzyme in *Sileneae*. Panel (A) shows the ancestral function of a single organellar aaRS enzyme that is imported by both mitochondria and chloroplasts. Panel (B) shows a duplication of the organellar enzyme with each paralog now specializing either on native plastid tRNAs or on cytosolic tRNAs imported into mitochondria that have lost the native tRNA gene. Panel (C) is a maximum likelihood tree of the complete organellar PheRS enzyme sequences from *A. thaliana* and *Sileneae* species with bootstrap values indicated on branches. To the right of the tree is an alignment of just the first 85 amino acids of the aaRS proteins. Right of the alignment are the targeting probabilities generated from the respective software program for each PheRS gene. All species shown have lost the corresponding mt-tRNA-Phe gene and are therefore underlined in bold font. The predicted transit peptide from the gene highlighted with a red dashed box was fused to GFP for colocalization analysis in panel D. (D) Transient expression of the predicted transit peptide from three different organellar PheRS enzymes in *N. benthamiana* epithelial cells. The amino acid sequence plus 10 upstream amino acids of the protein body were fused to GFP and co-transfected with an eqFP611-tagged transit peptide from a known mitochondrially localized protein (isovaleryl-CoA dehydrogenase).

### Mitochondrial localization of cytosolic aaRSs often happens prior to the loss of mt-tRNAs and can occur multiple times independently in a lineage

Phylogenetic comparisons indicated that the acquisition oftested from LysRS and TyrRS enzymes also resulted in GFP transit peptides by cytosolic aaRSs in *Sileneae* often occurred before the loss of the cognate mt-tRNA gene (Fig. 7). Only GlnRS (Fig. 4A), MetRS2 (Fig. 6C), and potentially TrpRS (Fig. 6B), showed a perfect match in the evolutionary timing of mt-tRNA loss and predicted cytosolic aaRS retargeting (Fig. 7). It should be noted that N-terminal extensions are present on cytosolic TrpRS genes in *S. latifolia* and *S. vulgaris* (both of which still retain a native mt-tRNA-Trp gene) (Fig. 6B), but they fell below the targeting prediction cutoff for mitochondrial localization. For the remaining cytosolic enzymes that gained predicted transit peptides (LysRS (Fig. 4A), TyrRS (Fig. 5A), and ProRS (Fig. 6A)), an N-terminal extension was also present in one or more species that still retained the mt-tRNA. Colocalization assays were performed in two such cases, confirming the ability of these extensions to target mitochondria (Figs. 4B and 5B). Because the organellar LysRS (Supp. Fig. 12), ProRS (Supp. Fig. 15), and TyrRS (Supp. Fig. 19) are still predicted to be mitochondrially localized, the apparent gain of mitochondrial targeting by the corresponding cytosolic aaRSs suggests that targeting of both enzymes prior to mt-tRNA loss is a widespread phenomenon in *Sileneae* (Fig. 7).

Although it was common for homologous transit peptides to be present in multiple species, there were also instances where transit peptides were gained independently multiple times for the same aaRS. A cytosolic ProRS in A. githago (Fig. 6A) and a cytosolic TyrRS in *S. vulgaris* (Fig. 5A) each had an N-terminal extension that was nonhomologous to the extensions found in other *Sileneae* species (i.e., no significant similarity with a blastn comparison at an e-value threshold of 0.1). In the case of cytosolic TyrRS in *S. vulgaris*, two different enzymes appear to have gained mitochondrial localization independently with two different N-terminal extensions (Fig. 5A). Representatives for each of these independently derived extensions were able to function as mitochondrial transit peptides in *N. benthamiana* (Fig. 5B). There were also cases where an N-terminal extension on an aaRS was unique to a single species. For example, we found a duplicate cytosolic AspRS gene in the nuclear genome assembly of *S. vulgaris* that was strongly predicted to be mitochondrially targeted, but no other *Sileneae* species appear to have gained mitochondrial targeting for AspRS (Supp. Fig. 4). In addition, there were multiple cases where a substantially truncated read or isoform resulted in predicted mitochondrial targeting (Supp. Table 2), but due to the length and low expression it was unclear if these products produce functional aaRSs or are just spurious sequencing or expression products. We therefore did not consider these AspRS and GluRS sequences to be likely cases where a cytosolic enzyme gained mitochondrial localization.

### Recently acquired transit peptides have no detectable homology with the transit peptides encoded by other genes in the genome

Transit peptides can evolve through duplication and transfer of transit peptides present on other existing genes (Liu et al., 2009; Wu et al., 2017). Therefore, we tested whether the transit peptides we identified in this study originated from other genes or evolved de novo from upstream regions. When putative transit peptides were searched against the nuclear genomes of each respective species, we found no cases where a transit peptide was donated to an aaRS from another protein. This is in agreement with studies that have found that de novo sequence evolution as the most common evolutionary mechanism in the transit peptide formation (Christian et al., 2020).

### Retargeting of cytosolic aaRSs to mitochondria may result in ancestrally dual-targeted organellar aaRSs now specializing exclusively in plastids

Predicting organelle-specific versus dual-targeted enzymes with purely in silico methods is difficult due to the shared characteristics of mitochondrial, plastid, and dual transit peptides. Nevertheless, we observed a decreased probability of aaRS enzymes being dual-targeted (and instead predicted to be only plastid localized) when a cytosolic enzyme gained a putative mitochondrial transit peptide. This pattern is consistent with expectations that functional replacement in the mitochondria will lead organellar aaRSs to function exclusively in the plastids.

The targeting of GlyRS enzymes presents an interesting situation in *A. thaliana* where both a cytosolic enzyme and a dual-targeted organellar enzyme are localized to the mitochondria (Fig. 7). In *Sileneae*, a putative transit peptide on the cytosolic GlyRS is also present, possibly being gained independently (Supp. Fig. 8). But unlike in *A. thaliana, Sileneae* species have lost the native mt-tRNA-Gly gene, suggesting a complete replacement of the bacterial Gly decoding system in *Sileneae* mitochondria. This functional replacement of tRNA/aaRS corresponds with a marked decreaseinthepredictedprobabilityofmitochondriallocalization of the organellar GlyRS enzyme resulting in an almost exclusively plastid-specific targeting prediction (Supp. Fig. 8).

Retargeting of cytosolic MetRS is also associated with changes in dual-targeting predictions for the organellar aaRSs. Although multiple organellar MetRS genes in *Sileneae* experienced only a marginal decrease in mitochondrial targeting prediction compared to *A. thaliana, S. vulgaris* had virtually no signal of mitochondrial localization (Supp. Fig. 13) and is the only species in the lineage that has lost both tRNA-Met genes (elongator Met and initiator fMet, Fig. 1). This observation raises the possibly that the loss of both tRNA genes has obviated the need for an organellar MetRS in *S. vulgaris* mitochondria, allowing the organellar MetRS to evolve exclusive plastid-targeting.

A similar reduction in mitochondrial targeting prediction was seen in organellar TrpRS enzymes, with the species that have lost the cognate mt-tRNA-Trp gene (and a predicted gain of mitochondrial targeting for the cytosolic TryRS enzyme), exhibiting organellar enzymes now predicted to be exclusively plastid localized (Supp. Fig. 18). Overall, plants appear to differ from systems such as nonbilaterian animals in which outright organellar aaRS loss has been observed in conjunction with replacement of their mitochondrial tRNA/aaRS system with cytosolic counterparts (Haen et al., 2010; Pett and Lavrov, 2015). In plants, the presence of plastids likely necessitates the retention of the enzyme. Whether there is selective pressure to specialize aaRS import to plastids once a cytosolic enzyme is localized to mitochondria or if the loss of dual targeting is just due to relaxed selection for function in mitochondria is unknown.

### Functional replacement of mt-tRNAs is not always associated with retargeting of cytosolic aaRSs and may sometimes require duplication and subfunctionalization of a dual-targeted enzyme

The repeated evolution of N-terminal transit peptides in *Sileneae* aaRSs (Fig. 4-6) supports a model of cytosolic retargeting as a key mechanism associated with changes in mt-tRNA content (Fig. 2A). However, there were also numerous examples where a mt-tRNA gene was lost (and functionally replaced by the import of a cytosolic tRNA) but there was no predicted change in cytosolic aaRS import (Fig. 7). For the cytosolic AsnRS, cytosolic CysRS, cytosolic HisRS, cytosolic PheRS, and cytosolic SerRS, organelle localization was not predicted by any of the software programs (Supp. Figs. 3, 5, 14, 16), and the length of the enzymes did not differ substantially from the corresponding *A. thaliana* ortholog(s) in alignments. As discussed above, it is also unlikely that cytosolic AspRS or GluRS gained mitochondrial targeting in *Sileneae*. Accordingly, the organellar aaRSs for Asn (Supp. Fig. 3), Asp (Supp. Fig. 4) Cys (Supp. Fig. 5), Glu (Supp. Fig. 7), His (Supp. Fig. 9), and Phe (Supp. Fig. 14) did retain predicted transit peptides for mitochondrial localization, suggesting that these organellar aaRSs are now charging the newly imported cytosolic tRNAs. The organellar SerRS retained a predicted transit peptide in *Sileneae*, but predictions were overwhelmingly for plastid localization, making it unclear if it still functions in the mitochondria (Supp. Fig. 16). In general, these examples appear to follow the model in which organellar aaRS now charge a novel (cytosolic) tRNA substrate (Fig. 2B). However, mitochondrial targeting of aaRSs (and proteins more generally) is not always based on identifiable N-terminal transit peptides (Duchêne et al., 2005; Dudek et al., 2013; Reinbothe et al., 2021), so it is possible that additional cytosolic aaRSs are imported into mitochondria but were not detected in this analysis.

Nevertheless, some organellar aaRSs are known to be less discriminating than bacterial or cytosolic counterparts (Salinas-Giegé et al., 2015), so it is possible that these organellar enzymes are inherently permissive enzymes capable of charging newly imported, cytosolic tRNAs (also see section on identity elements below). Alternatively, adaptive amino acid substitutions in an organellar enzyme could facilitate cytosolic tRNA recognition. This scenario of aaRS adaptation raises the possibility of pleiotropic effects on plastid translation, as a dual targeted organellar aaRS would have to adapt to charge cytosolic tRNAs but also maintain aminoacylation function in the plastid with bacterial-like, plastid tRNAs.

PheRS presented a unique case where an organellar enzyme appears to be charging imported, nuclear-encoded tRNAs, but the ancestrally a dual-targeted enzyme has undergone duplication and subfunctionalization in *Sileneae* such that one copy is specifically plastid-localized (Fig. 8A-B). In *A. thaliana*, only a single organellar PheRS has been found, and fusion of that transit peptide to GFP resulted in dual localization to both organelles (Fig. 8C). This suggests that the organellar *A. thaliana* PheRS enzyme can charge native plastid tRNAs as well as imported tRNA-Phe (*A. thaliana* has also lost mt-tRNA-Phe). However, the enzymatic coevolutionary response to losing this mt-tRNA may be sustainably different in *Sileneae* as there has been a gene duplication event in the organellar PheRS gene family where one of the PheRS paralogs has a stronger prediction for mitochondrial targeting than plastid targeting, and the inverse is true for the other paralog (Fig. 8C). Accordingly, the predicted mitochondrial transit peptide for PheRS in *S. conica* showed strong mitochondrial, and not plastid, targeting in colocalization assays (Fig. 8D [76028]. Similarly, the predicted plastid transit peptide for *S. conica* PheRS showed primarily plastid localization and only very weak mitochondrial localization in these assays (Fig. 8D). The duplication and apparent subfunctionalization of organellar PheRS may have been necessary because of constraints in cellular trafficking. The mt-tRNA-Phe is the only mt-tRNA lost three times independently in this angiosperm dataset, yet there is no evidence of cytosolic PheRS gaining mitochondrial import in any of these lineages. Notably, cytosolic PheRS is the only aaRS composed of two heterodimers with essential α- and β-subunits (Safro et al., 2013). The import of both subunits and successful assembly of the dimer is presumably essential for aminoacylation inside the mitochondrial matrix, thus requiring the almost simultaneous acquisition of a targeting peptide on both subunits for functional replacement. This import requirement may pose an unusually difficult “two-body problem” to functionally replace the organellar PheRS with its multi-subunit cytosolic counterpart. Similarly, mitochondrial PheRS has never been replaced in animals despite mt-tRNA-Phe being lost at least three times in that branch of eukaryotes (Pett and Lavrov, 2015). We hypothesize that the mitochondrial specialization of one of these organellar-targeted paralogs in *Sileneae* may indicates adaptation to recognize the novel, cytosolic tRNAs – an enzymatic change that could interfere with the charging of plastid tRNAs and necessitate two subfunctionalized enzymes.

### Shared discriminator bases between nuclear-encoded and native mitochondrial tRNAs may facilitate organellar aaRS recognition of both tRNA classes

Our results indicate that roughly half of the examples support each of the two very different routes to the replacement of the bacterial aaRS/tRNA system in plant mitochondria (permissive aaRSs and redundant aaRS import; Fig. 2). These findings may offer insight into each enzyme’s activity and address a striking contrast encountered in aaRS evolution. On one hand, aaRSs have successfully undergone horizontal gene transfer across some of the deepest splits in tree of life without disrupting their function (Doolittle and Handy, 1998; Brindefalk et al., 2007). On the other hand, aaRSs are also highly discriminating enzymes. Even within mitochondrial translation systems, there are multiple examples of single nucleotide substitutions in mt-tRNAs resulting in severe reductions in aminoacylation (Yarham et al., 2010). In one described case of aaRS/tRNA incompatibility in Drosophila, a single amino acid polymorphism in the mitochondrial TyrRS negatively interacted with a nucleotide polymorphism in mt-tRNA-Tyr to produce a diseased phenotype of delayed development and reduced fecundity (Meiklejohn et al., 2013). The replacement of a mt-tRNA with a cytosolic tRNA represents a far more radical change in substrates, and raises the following question: Are there specific features of aaRS-tRNA relationships that make them more or less likely to follow one of the two alternative evolutionary paths to functional replacement?

One possibility is that organellar aaRSs are predisposed to recognize cytosolic tRNAs when key identity elements necessary for recognition and charging happen to be shared between cytosolic tRNAs and native mt-tRNAs (Fig. 2B). In contrast, retargeting of cytosolic aaRSs might be favored when cytosolic tRNAs and mt-tRNAs differ in key identify elements (Fig. 2A). The positions of identity elements vary among tRNA families, but there are some common themes, including the near-universal role of the discriminator base, i.e., the nucleotide at the 3′ end of each tRNA prior to the addition of the CCA tail (Giegé et al., 1998). Therefore, to investigate how differences in identity elements between mitochondrial and cytosolic tRNAs might affect aaRS recognition in cases of mt-tRNA gene loss and functional replacement, we compared typical angiosperm discriminator bases in cytosolic, mitochondrial, and plastid tRNAs (Table 1).

**Table 1.**
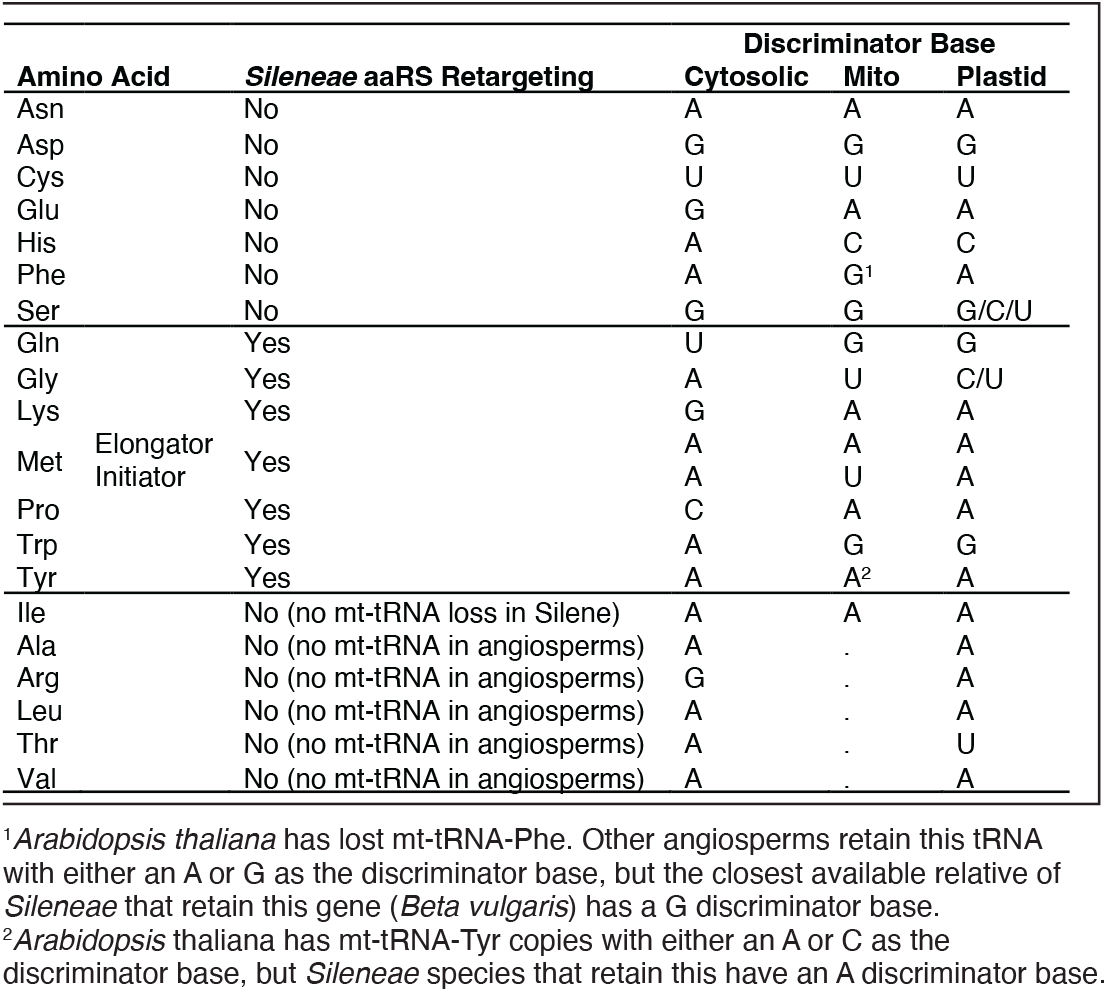
Discriminator bases in *Arabidopsis thaliana* cytosolic, mitochondrial, and plastid tRNAs as obtained the PlantRNA database (Cognat et al., 2021). Amino acids are organized into groups based on the evolutionary history of mt-tRNA gene loss and predicted cytosolic aaRS retargeting in *Sileneae*.

There are seven cytosolic aaRSs that are predicted to be targeted to the mitochondria in *Sileneae* in association with loss and functional replacement of cognate mt-tRNA genes: GlnRS, GlyRS, LysRS, MetRS, ProRS, TrpRS, and TyrRS (Fig. 7). In six of these seven cases, angiosperm mitochondrial and cytosolic tRNAs typically differ in their discriminator bases (Table 1; note that elongator tRNA-Met genes share the same discriminator base, but MetRS must also charge initiator tRNA-Met, which has different discriminator bases in its mt and cytosolic versions). The only exception among these seven cases is tRNA-Tyr, but the mitochondrial and cytosolic versions of this tRNA differ in its other key identity element – the paired bases at the end of its acceptor stem (Tsunoda et al., 2007). Bacterial (including plant mt- and plastid) tRNA-Tyr generally has a G1-C72 base-pair, whereas the eukaryotic (cytosolic) counterpart has a C1-G72 pair (Cognat et al., 2021). Even though vertebrate mitochondrial TyrRSs have apparently lost their ability to distinguish between these alternative identity elements (Bonnefond et al., 2005), the organellar TyrRS in plants is independently derived from a cyanobacterial-like (presumably plastid) lineage (Duchêne et al., 2005; Brandao and Silva-Filho, 2011). Thus, these differences in putative identity elements may be one reason why retargeting of cytosolic GlnRS, GlyRS, LysRS, MetRS, ProRS, TrpRS, and TyrRS was necessary to facilitate the import and function of these cytosolic tRNAs into the mitochondria.

For seven other amino acids, the loss of a mt-tRNA did not appear to be associated with the retargeting of the corresponding cytosolic aaRS: AsnRS, AspRS, CysRS, GluRS, HisRS, PheRS, and SerRS (Fig. 7). Therefore, in these cases, it appears that the native organellar aaRS retains mitochondrial localization and now charges cytosolic tRNAs that are newly imported into the mitochondria, although it is possible that cytosolic aaRS retargeting has occurred but is not detectable with in silico predication algorithms. In four of these seven cases, the same discriminator base is typically used in plant mitochondrial and cytosolic tRNAs: tRNA-Asn, tRNA-Asp, tRNA-Cys, and tRNA-Ser (Table 1), perhaps contributing to the ability of organellar aaRSs to charge cytosolic-like tRNAs.

Even though there are differences between mitochondrial and cytosolic discriminator bases in the remaining three cases (tRNA-Glu, tRNA-His, and tRNA-Phe), there are reasons to believe that these differences may not interfere with aaRS specificity. In particular, tRNA-Glu is one of only two examples (tRNA-Thr being the other) where the discriminator base has not been found to act as an identity element in bacterial-like tRNAs (Giegé et al., 1998). In the case of tRNA-Phe, the native mt-tRNA genes found across angiosperms exhibit variation in the discriminator base and can have either an A or a G at this position. Therefore, the plant organellar PheRS may have already evolved to recognize either of these two alternative nucleotides, which would be consistent with the permissive nature of mt PheRS in humans (Klipcan et al., 2012; Salinas-Giegé et al., 2015). HisRS has an exceptionally complex evolutionary history (Duchêne et al., 2005; Ardell and Andersson, 2006; Brindefalk et al., 2007). Most bacterial and archaeal HisRS enzymes have a conserved Gln residue that directly interacts with the C discriminator base in prokaryotic tRNA-His and likely determines specificity (Ardell and Andersson, 2006; Lee et al., 2017). In contrast, eukaryotic cytosolic aaRSs lack this Gln residue, and cytosolic tRNA-His typically has an A nucleotide at the discriminator base position (Giegé et al., 1998; Lee et al., 2017). Many animals and fungi only have a single HisRS, which is capable of charging both mt and cytosolic tRNA-His (Lee et al., 2017). Plants, however, have a distinct organellar HisRS. Even though this plant organellar HisRS appears to be of archaeal origin (Duchêne et al., 2005), it has lost the conserved Gln residue typically present in prokaryotic HisRSs and has converged on a Met-Thr motif at this position that is also found in the main family of eukaryotic HisRSs (Lee et al., 2017). Therefore, the plant organellar HisRS may be more permissive in charging tRNAs with either discriminator base like that of the sole HisRS found in many eukaryotes. This is consistent with a more general observation across eukaryotes that mt aaRSs often evolve to be more permissive in tRNA charging (Kumazawa et al., 1991; Bonnefond et al., 2005; Fender et al., 2006).

Overall, these comparisons of discriminator bases provide suggestive evidence that the extent of similarity in identity elements between mt and cytosolic tRNAs may have shaped the evolutionary pathways associated with mt-tRNA gene loss and functional replacement (Fig. 2). In cases where mt and cytosolic tRNAs are sufficiently similar in identity elements, the organellar aaRSs may be able to persist in the mitochondria and charge newly imported cytosolic tRNAs without major changes in sequence. However, given that identity elements can be found in many positions other than the discriminator base and that their locations vary idiosyncratically among tRNA families (Giege et al., 1998), a more detailed analysis of contact interfaces between aaRSs and tRNAs, as well as functional charging assays, will be needed to fully address this question.

### The chicken-or-the-egg problem of mt-tRNA replacement

One longstanding question related to mt-tRNA replacement in plants is whether tRNA or aaRS import happens first, as it has been assumed that the import of one without the other would be nonfunctional in translation or even toxic (Small et al., 1999). Our results provide evidence for two different scenarios that likely facilitate the loss of mt-tRNAs. As described above, it is possible that enzymatic flexibility and/ or shared identity elements on some mt- and cytosolic tRNAs have resulted in permissive aaRS/tRNA interactions ensuing the charging of nuclear-encoded tRNAs by organellar enzymes (Fig. 2B). Furthermore, recent work to detect tRNA import in *Sileneae* found cases of redundant import of a cytosolic tRNA prior to the loss of the mt-tRNA gene for tRNA-Asn, tRNA-Glu, and tRNA-His (Warren et al., 2021). The results from the present study suggest that these corresponding organellar aaRSs are capable of charging all three of these cytosolic tRNAs (Supp. Fig. 3, Supp. Fig. 7, Supp. Fig. 9), setting up a “tRNA-first” transition state. Once both tRNAs are functional within the mitochondria, it becomes easy to envision a scenario where an inactivating mutation in the mt-tRNA gene makes the system wholly dependent on the cytosolic tRNA.

The second potential transition state involves the initial evolution of cytosolic aaRS import (Fig. 2A) with little or no cognate tRNA import. There is some indication that this state can occur, as we previously found that cytosolic tRNA-Tyr was very depleted in *S. vulgaris* mitochondria (Warren et al., 2021), yet here we found evidence for the import of two copies of the cytosolic TyrRS enzyme in the same species (Fig. 5). Therefore, it is possible that these imported aaRSs have a function other than aminoacylation or have some activity on mitochondrial tRNAs. More generally, we found evidence for multiple aaRSs (Lys, Pro, and Tyr) that cytosolic- and organellar-like enzymes could both be present in the mitochondria and that gain of cytosolic aaRS import preceded loss of the corresponding mt-tRNA gene. Such patterns are expected under an “aaRS-first” model, but they do not offer conclusive support especially because it is difficult to ever demonstrate that mitochondrial import of a particular cytosolic tRNA is completely absent. In an aaRS-first scenario, the eventual import of the cytosolic tRNA would then give rise to an intermediate state of mitochondrial translation where both the organellar system (a bacterial-like tRNA/aaRS) and a cytosolic system (eukaryotic-like tRNA/ aaRS) are cofunctional in mitochondria. Such a situation exists in *A. thaliana* where both imported tRNA-Gly and mt-tRNA-Gly are necessary for translation (Salinas et al., 2005). The presence of both imported and native tRNAs that decode the same amino acid (but different codons) is mirrored by the import of both an organellar and cytosolic GlyRS (Fig. 7). The bacterial-like GlyRS was found to effectively aminoacylate both tRNA counterparts, whereas the cytosolic GlyRS had poor activity with a mt-tRNA-Gly substrate (Duchêne and Marechal-Drouard, 2001). It would be interesting to determine whether similar scenarios exist in *Sileneae* where a cytosolic aaRS has cross-functionality in charging both tRNAs.

In summary, the repeated loss and functional replacement of mt-tRNA genes in plants does not appear to involve a single order of evolutionary events or even a single eventual end-state. In some cases, early retargeting of aaRSs to the mitochondria is likely key to the process, but in others, import of cytosolic tRNAs clearly occurs first. Indeed, the replacement of mt-tRNA genes may sometimes follow a “tRNA-only” model, as we have shown that full loss of mt-tRNA genes can occur without any apparent retargeting of cytosolic aaRSs. Which of these trajectories is taken is unlikely to be entirely random. Instead, the evolutionary pathway may be influenced by the molecular and enzymatic features of tRNA/ aaRS interactions, such as sharing of identity elements between cytosolic and mitochondrial tRNAs or constraints on import imposed by a multisubunit enzyme (PheRS). In addition, this evolutionary process may be shaped by the distinctive tripartite translation system in plants, which requires that plastid functions be preserved even during periods of dynamic change in mitochondrial translation.

## Materials and Methods

### Tissue generation and growth conditions

Tissue generation, RNA extraction, and Iso-Seq library construction for *S. noctiflora* were done in a previously described study (Williams et al., 2021), while data for the other four *Sileneae* species were newly generated for this study. The following seed collections or accessions were used: A. githago Kew Gardens Millennium Seed Bank (0053084), *S. vulgaris* S9L (Sloan et al., 2012a), *S. latifolia* UK2600 (from the line originally used for mitogenome sequencing in (Sloan et al., 2010)), and *S. conica* ABR (Sloan et al., 2012b). Seeds were germinated in small plastic pots with Plantorium Greenhouse brand potting soil in a growth chamber at 23 °C with a light setting of 8-hour light/16-hour dark at 100 *µ*E m-1 s-1. One week after germination, chamber settings were modified to promote flowering (“long-day” conditions) with 16-hour light/8-hour dark.

### RNA extraction and Iso-Seq library construction

RNA was extracted from *A. githago* (hermaphrodite), *S. conica* (hermaphrodite), *S. latifolia* (male), and *S. vulgaris* (male-fertile hermaphrodite) with a Qiagen RNeasy Plant Mini Kit, using RLT buffer with 10 *µ*l beta-mercaptoethanol. RNA was DNase treated with a Qiagen RNase-Free DNase Set. Separate RNA extractions were performed on leaf tissue and an immature flower sample (∼5 days post flower development) for *A. githago, S. vulgaris, and S. latifolia*. Two different tissues were used to increase detection of diverse transcripts, but the two RNA samples were pooled equally by mass for each species prior to library construction, so individual reads cannot be assigned to leaf or floral tissues. Only leaf tissue was used for *S. conica* as the individual had not yet begun flowering at the time of RNA extraction. Both tissue types were harvested at 4 weeks post-germination, and RNA integrity and purity were checked on a TapeStation 2200 and a Nanodrop 2000.

Iso-Seq library construction and sequencing were performed at the Arizona Genomics Institute. Library construction was done using PacBio’s SMRTbell Express Template Prep Kit 2.0. The four libraries were barcoded and pooled. The multiplexed pool was sequenced with a PacBio Sequel II platform on two SMRT Cells using a Sequencing Primer V4, Sequel II Bind Kit 2.0, Internal Control 1.0, and Sequel II Sequencing Kit 2.0. Raw movie files were processed to generate circular consensus sequences (CCSs) using PacBio’s SMRT Link v9.0.0.92188 software (Pacific Biosciences 2020). Demultiplexing was performed with lima v2.0.0 and the --isoseq option. Full-length non-chimeric (FLNC) sequences were generated with the refine command and the --require_polya option in the IsoSeq3 (v3.4.0) pipeline. Clustering of FLNCs into isoforms was then performed with the cluster command in IsoSeq3 with the --use-qvs option. The two SMRT Cells produced similar outputs with 5.8M and 5.9M raw reads, which resulted in 3.9M CCSs for each cell (3.5M and 3.4M retained after demultiplexing). The results of demultiplexing, FLNC filtering, and clustering are shown in Supp. Table 3.

### Extraction of aaRS transcript sequences

*Arabidopsis* aaRS genes were identified from published sources (Duchêne et al., 2005; Warren and Sloan, 2020) and the corresponding protein sequences were obtained from the Araport11 genome annotation (201606 release). Homologs from the high-quality (HQ) clustered isoforms from each species were identified with a custom Perl script (iso-seq_blast_pipeline.pl available at GitHub: https://github.com/warrenjessica/Iso-Seq_scripts) that performed a tBLASTn search with each *Arabidopsis* aaRS sequence, requiring a minimum sequence identity of 50% and a minimum query length coverage of 50%. All HQ clusters that satisfied these criteria were retained by setting the --min_read parameter to 2 (the IsoSeq3 clustering step already excludes singleton transcripts).

### Transcript processing and targeting prediction

The longest ORF was extracted from each aaRS transcript using the EMBOSS v. 6.6.0 (Rice et al., 2000) getorf program with the options: -minsize 75 -find 1. Many Iso-Seq transcripts differed in length by only a few nucleotides in UTRs but resulted in identical ORFs. Therefore, all identical ORFs were collapsed for downstream targeting and phylogenetic analysis. Collapsed ORFs were translated into protein coding sequences for localization analysis. TargetP v.2.0 (Almagro Armenteros et al., 2019), LOCALIZER v.1.0.4 (Sperschneider et al., 2017), and Predotar v.1.04 (Small et al., 2004) were each used to predict targeting probabilities of each coding sequence. All programs were run with the plant option.

### Determination of gene copy number and genome assembly scanning for undetected genes

Very similar transcripts can be the product of different genes, alleles, or sequencing errors. In order to infer the number of unique genes for each related set of transcripts in a species, CD-HIT-EST v. 4.8.1 (Fu et al., 2012) was used to further cluster transcripts into groups. For this clustering step, sequences were first aligned with MAFFT v. 7.245 (Katoh and Standley, 2013) with default settings and trimmed by eye to remove terminal sequence ends with gaps and N-terminal extensions that were not present on all sequences. Any two sequences in which the coding region shared greater than 98% sequence similarity were collapsed into a single gene cluster (CD-HIT-EST options -c 0.98 -n 5 -d 0). Each cluster of transcripts was considered a single gene, and the transcript with the highest expression and longest length was retained as the representative sequence for the gene. To check for the possibility that a cytosolic gene had gained a transit peptide but was undetected in Iso-Seq data (due to low expression or representation in the sequencing library), all cytosolic genes that appeared to lack transit peptides were checked for immediately upstream start codons in the corresponding nuclear genome assembly (Warren et al., 2021). Representative transcripts from each gene cluster were translated and BLASTed (tblastn) against the nuclear assembly, and scaffolds with a hit to the first exon of the protein were extracted and analyzed with the ExPASy Translate tool (Artimo et al., 2012). The ORF found in the genome assembly was then compared to the ORF generated from the transcript and inspected for length differences. If an upstream Met was present, the upstream sequence was appended to the rest of the gene and re-run through the targeting prediction software described above. Occasionally, when BLASTing cytosolic aaRS proteins to nuclear assemblies, additional genes were discovered that were entirely absent from the Iso-Seq data (genes marked with ** in Supp. Figs. 3,4,13 and 19). In these cases, the region that aligned to the first exon of the expressed paralog was used for phylogenetic and targeting analysis.

### Sequence alignment and maximum likelihood phylogenetic analysis

After clustering transcripts by sequence similarity (see above), the coding region of the longest transcript for each gene was retained for phylogenetic analysis. If two or more transcripts were tied for the longest length, the one with higher expression level was used. Retained sequences for each aaRS gene family were aligned using MAFFT v. 7.245 (Katoh and Standley, 2013) with default settings. Sequences were trimmed by eye to remove poorly aligned regions, and maximum likelihood trees were produced using RAxML v.8.2.12 (Stamatakis, 2014) with a GTRGAMMA model and rapid bootstrap analysis with a 100 replicates. Sequence alignments for Figs. 4-6, 8 and Supp. Fig. 8 were generated in Geneious (Geneious Prime 2022.2.2, https://www.geneious.com) (parameters: geneious alignment, global with free end gaps, Blosum62) with the full amino acid sequence. A window of the first ∼100 aligned N-terminal amino acids from the alignment was loaded with the corresponding trees into the R package ggtree (Yu, 2020) to generate alignment figures.

### Transient expression of transit peptides and colocalization assays in N. benthamiana epithelial cells

Constructs were made from putative transit peptides predicted from TargetP v.2.0 (Almagro Armenteros et al., 2019). Each transit peptide plus the following 30 bp (10 amino acids) was placed between the attLR1 (5’) and attLR2 (3’) Gateway cloning sites. The desired constructs were synthesized and cloned into pUC57 (Ampr) using EcoRI and BamHI restriction sites by GenScript, transferred into the constitutive plant destination vector pK7FWG2 (bacterial Specr/plant Kanr) (Karimi et al., 2002), which contains a C-terminal GFP fusion, using Gateway LR Clonase II Enzyme Mix, and transformed into *E. coli* DH5a. Two colonies were selected for each construct, DNA was purified using the GeneJet Plasmid Miniprep Kit (Thermo Scientific) and verified by full-length plasmid sequencing (Plasmidsaurus). The putative transit peptides and following 10 amino acids were confirmed to be in-frame with the C-terminal GFP fusion protein by sequence alignment. Positive clones were used to transform electrocompetent Agrobacterium C58C1-RifR (also known as GV3101::pMP90, (Hellens et al., 2000)), colonies were selected on Rif/Spec/Gent (50 *µ*g/mL each) and confirmed by PCR using primers directed to the 5’ (Cam35S promoter) and 3’ (GFP) regions flanking the constructs.

*Agrobacterium* transient transformation of *N. benthamiana* leaves was done using the method of Mangano et al. (2014), but scaled up to accommodate *N. benthamiana* instead of *Arabidopsis* leaves. The species N. benthamiana was used for transformation because it does not have a hypersensitive response to *Agrobacterium* at the infiltration site.

Leaf samples were imaged after 48 hr on a Nikon A1-NiE confocal microscope equipped with a CFI Plan Apo VC 60 XC WI objective. GFP, eqFP611, and chlorophyll were excited and collected sequentially using the following excitation/emissions wavelengths: 488 nm / 525/50 nm (GFP), 561 nm / 595/50 nm (red fluorescent protein eqFP611), 640 nm / 700 (663 – 738) nm (chlorophylls). Imaging was done using Nikon NIS-Elements 5.21.03 (Build 1489), and image analysis was performed using Nikon NIS-Elements 5.41.01 (Build 1709). Maximum Intensity Projections in Z were produced after using the Align Current ND Document (settings: Align to Previous Frame, The intersection of moved images, Process the entire image), and 500 pixel 500 pixel (103.56 *µ*M × 103.56 *µ*M) cropped images were created from each projection for figures.

## Data Availability

The CCSs from each Iso-Seq library are available via the NCBI Sequence Read Archive (SRA) under BioProject PRJNA799780. Trimmed and untrimmed alignments for final aaRS sequences,as well as raw microscopy image files,can be found on Dryad at https://doi.org/10.5061/dryad.0k6djhb20.

## Acknowledgements

We thank Salah Abdel-Ghany and Rachael DeTar for help in designing the cloning strategy used for colocalization assays. This work was supported by grants from the National Science Foundation (MCB-2048407 and MCB-1933590).

## References

Almagro Armenteros, J.J., Salvatore, M., Emanuelsson, O., Winther, O., von Heijne, G., Elofsson, A., and Nielsen, H. (2019). Detecting sequence signals in targeting peptides using deep learning. Life Sci Alliance 2.

Ardell, D.H., and Andersson, S.G. (2006). TFAM detects co-evolution of tRNA identity rules with lateral transfer of histidyl-tRNA synthetase. Nucleic Acids Res 34, 893–904.

Artimo, P., Jonnalagedda, M., Arnold, K., Baratin, D., Csardi, G., de Castro, E., Duvaud, S., Flegel, V., Fortier, A., Gasteiger, E., Grosdidier, A., Hernandez, C., Ioannidis, V., Kuznetsov, D., Liechti, R., Moretti, S., Mostaguir, K., Redaschi, N., Rossier, G., Xenarios, I., and Stockinger, H. (2012). ExPASy: SIB bioinformatics resource portal. Nucleic Acids Res 40, W597–603.

Berglund, A.K., Spanning, E., Biverstahl, H., Maddalo, G., Tellgren-Roth, C., Maler, L., and Glaser, E. (2009). Dual targeting to mitochondria and chloroplasts: characterization of Thr-tRNA synthetase targeting peptide. Mol Plant 2, 1298–1309.

Bonnefond, L., Frugier, M., Giegé, R., and Rudinger-Thirion, J. (2005). Human mitochondrial TyrRS disobeys the tyrosine identity rules. RNA 11, 558–562.

Boore, J.L. (1999). Animal mitochondrial genomes. Nucleic Acids Res 27, 1767–1780.

Brandao, M.M., and Silva-Filho, M.C. (2011). Evolutionary history of Arabidopsis thaliana aminoacyl-tRNA synthetase dual-targeted proteins. Mol Biol Evol 28, 79–85.

Brindefalk, B., Viklund, J., Larsson, D., Thollesson, M., and Andersson, S.G. (2007). Origin and evolution of the mitochondrial aminoacyl-tRNA synthetases. Mol Biol Evol 24, 743–756.

Bruce, B.D. (2001). The paradox of plastid transit peptides: conservation of function despite divergence in primary structure. Biochim Biophys Acta 1541, 2–21.

Christian, R.W., Hewitt, S.L., Nelson, G., Roalson, E.H., and Dhingra, A. (2020). Plastid transit peptides-where do they come from and where do they all belong? Multi-genome and pan-genomic assessment of chloroplast transit peptide evolution. PeerJ 8, e9772.

Cognat, V., Pawlak, G., Pflieger, D., and Drouard, L. (2021). PlantRNA 2.0: an updated database dedicated to tRNAs of photosynthetic eukaryotes. bioRxiv, 2021.2012.2021.473619.

Delage, L., Dietrich, A., Cosset, A., and Marechal-Drouard, L. (2003). In vitro import of a nuclearly encoded tRNA into mitochondria of Solanum tuberosum. Mol Cell Biol 23, 4000–4012.

Doolittle, R.F., and Handy, J. (1998). Evolutionary anomalies among the aminoacyl-tRNA synthetases. Curr Opin Genet Dev 8, 630–636.

Duchêne, A.M., and Marechal-Drouard, L. (2001). The chloroplast-derived trnW and trnM-e genes are not expressed in Arabidopsis mitochondria. Biochem Biophys Res Commun 285, 1213–1216.

Duchêne, A.M., Pujol, C., and Marechal-Drouard, L. (2009). Import of tRNAs and aminoacyl-tRNA synthetases into mitochondria. Curr Genet 55, 1–18.

Duchêne, A.M., Giritch, A., Hoffmann, B., Cognat, V., Lancelin, D., Peeters, N.M., Zaepfel, M., Marechal-Drouard, L., and Small, I.D. (2005). Dual targeting is the rule for organellar aminoacyl-tRNA synthetases in Arabidopsis thaliana. P Natl Acad Sci USA 102, 16484–16489.

Dudek, J., Rehling, P., and van der Laan, M. (2013). Mitochondrial protein import: common principles and physiological networks. Biochim Biophys Acta 1833, 274–285.

Fender, A., Sauter, C., Messmer, M., Pütz, J., Giegé, R., Florentz, C., and Sissler, M. (2006). Loss of a primordial identity element for a mammalian mitochondrial aminoacylation system. Journal of Biological Chemistry 281, 15980–15986.

Fu, L., Niu, B., Zhu, Z., Wu, S., and Li, W. (2012). CD-HIT: accelerated for clustering the next-generation sequencing data. Bioinformatics 28, 3150–3152.

Ge, C., Spanning, E., Glaser, E., and Wieslander, A. (2014). Import determinants of organelle-specific and dual targeting peptides of mitochondria and chloroplasts in Arabidopsis thaliana. Mol Plant 7, 121–136.

Ghifari, A.S., Gill-Hille, M., and Murcha, M.W. (2018). Plant mitochondrial protein import: the ins and outs. Biochem J 475, 2191–2208.

Giannakis, K., Arrowsmith, S.J., Richards, L., Gasparini, S., Chustecki, J.M., Royrvik, E.C., and Johnston, I.G. (2022). Evolutionary inference across eukaryotes identifies universal features shaping organelle gene retention. Cell Syst 13, 874–884 e875.

Giegé, R., Sissler, M., and Florentz, C. (1998). Universal rules and idiosyncratic features in tRNA identity. Nucleic Acids Res 26, 5017–5035.

Haen, K.M., Pett, W., and Lavrov, D.V. (2010). Parallel Loss of Nuclear-Encoded Mitochondrial Aminoacyl-tRNA Synthetases and mtDNA-Encoded tRNAs in Cnidaria. Mol Biol Evol 27, 2216–2219.

Hellens, R., Mullineaux, P., and Klee, H. (2000). Technical Focus:a guide to Agrobacterium binary Ti vectors. Trends Plant Sci 5, 446–451.

Huang, C.Y., Ayliffe, M.A., and Timmis, J.N. (2003). Direct measurement of the transfer rate of chloroplast DNA into the nucleus. Nature 422, 72–76.

Huang, S., Taylor, N.L., Whelan, J., and Millar, A.H. (2009). Refining the definition of plant mitochondrial presequences through analysis of sorting signals, N-terminal modifications, and cleavage motifs. Plant Physiol 150, 1272–1285.

Karimi, M., Inze, D., and Depicker, A. (2002). GATEWAY vectors for Agrobacterium-mediated plant transformation. Trends Plant Sci 7, 193–195.

Katoh, K., and Standley, D.M. (2013). MAFFT Multiple Sequence Alignment Software Version 7: Improvements in Performance and Usability. Mol Biol Evol 30, 772–780.

Klipcan, L., Moor, N., Finarov, I., Kessler, N., Sukhanova, M., and Safro, M.G. (2012). Crystal structure of human mitochondrial PheRS complexed with tRNAPhe in the active “open” state. Journal of Molecular Biology 415, 527–537.

Krasovec, M., Chester, M., Ridout, K., and Filatov, D.A. (2018). The Mutation Rate and the Age of the Sex Chromosomes in Silene latifolia. Curr Biol 28, 1832–1838 e1834.

Kumazawa, Y., Himeno, H., Miura, K.-i., and Watanabe, K. (1991). Unilateral aminoacylation specificity between bovine mitochondria and eubacteria. Journal of Biochemistry 109, 421–427.

Kunze, M., and Berger, J. (2015). The similarity between N-terminal targeting signals for protein import into different organelles and its evolutionary relevance. Front Physiol 6, 259.

Lee, D.W., Kim, J.K., Lee, S., Choi, S., Kim, S., and Hwang, I. (2008). Arabidopsis nuclear-encoded plastid transit peptides contain multiple sequence subgroups with distinctive chloroplast-targeting sequence motifs. Plant Cell 20, 1603–1622.

Lee, Y.-H., Chang, C.-P., Cheng, Y.-J., Kuo, Y.-Y., Lin, Y.-S., and Wang, C.-C. (2017). Evolutionary gain of highly divergent tRNA specificities by two isoforms of human histidyl-tRNA synthetase. Cellular and Molecular Life Sciences 74, 2663–2677.

Liu, S.L., Zhuang, Y., Zhang, P., and Adams, K.L. (2009). Comparative Analysis of Structural Diversity and Sequence Evolution in Plant Mitochondrial Genes Transferred to the Nucleus. Mol Biol Evol 26, 875–891.

Lynch, M. (2007). The origins of genome architecture. (Sunderland, Mass.: Sinauer Associates).

Mangano, S., Gonzalez, C.D., and Petruccelli, S. (2014). Agrobacterium tumefaciens-mediated transient transformation of Arabidopsis thaliana leaves. Methods Mol Biol 1062, 165–173.

Meiklejohn, C.D., Holmbeck, M.A., Siddiq, M.A., Abt, D.N., Rand, D.M., and Montooth, K.L. (2013). An Incompatibility between a Mitochondrial tRNA and Its Nuclear-Encoded tRNA Synthetase Compromises Development and Fitness in Drosophila. Plos Genet 9.

Michaud, M., Cognat, V., Duchêne, A.M., and Marechal-Drouard, L. (2011). A global picture of tRNA genes in plant genomes. Plant J 66, 80–93.

Mireau, H., Lancelin, D., and Small, I.D. (1996). The same Arabidopsis gene encodes both cytosolic and mitochondrial alanyl-tRNA synthetases. Plant Cell 8, 1027–1039.

Murcha, M.W., Kmiec, B., Kubiszewski-Jakubiak, S., Teixeira, P.F., Glaser, E., and Whelan, J. (2014). Protein import into plant mitochondria: signals, machinery, processing, and regulation. J Exp Bot 65, 6301–6335.

Peeters, N., and Small, I. (2001). Dual targeting to mitochondria and chloroplasts. Biochim Biophys Acta 1541, 54–63.

Petersen, G., Cuenca, A., Moller, I.M., and Seberg, O. (2015). Massive gene loss in mistletoe (Viscum, Viscaceae) mitochondria. Sci Rep 5, 17588.

Pett, W., and Lavrov, D.V. (2015). Cytonuclear Interactions in the Evolution of Animal Mitochondrial tRNA Metabolism. Genome Biol Evol 7, 2089–2101.

Pujol, C., Marechal-Drouard, L., and Duchêne, A.M. (2007). How can organellar protein N-terminal sequences be dual targeting signals? In silico analysis and mutagenesis approach. J Mol Biol 369, 356–367.

Reinbothe, S., Rossig, C., Gray, J., Rustgi, S., von Wettstein, D., Reinbothe, C., and Rassow, J. (2021). tRNA-Dependent Import of a Transit Sequence-Less Aminoacyl-tRNA Synthetase (LeuRS2) into the Mitochondria of Arabidopsis. Int J Mol Sci 22.

Rice, P., Longden, I., and Bleasby, A. (2000). EMBOSS: the European Molecular Biology Open Software Suite. Trends Genet 16, 276–277.

Rubio Gomez, M.A., and Ibba, M. (2020). Aminoacyl-tRNA synthetases. RNA 26, 910–936.

Safro, G., Moor, N., and Lavrik, O. (2013). Phenylalanyl-tRNA Synthetases. In: Madame Curie Bioscience Database [Internet].

Salinas, T., Schaeffer, C., Marechal-Drouard, L., and Duchêne, A.M. (2005). Sequence dependence of tRNA(Gly) import into tobacco mitochondria. Biochimie 87, 863–872.

Salinas-Giegé, T., Giegé, R., and Giegé, P. (2015). tRNA biology in mitochondria. Int J Mol Sci 16, 4518–4559.

Schmidt, O., Pfanner, N., and Meisinger, C. (2010). Mitochondrial protein import: from proteomics to functional mechanisms. Nat Rev Mol Cell Biol 11, 655–667.

Sloan, D.B., Alverson, A.J., Storchova, H., Palmer, J.D., and Taylor, D.R. (2010). Extensive loss of translational genes in the structurally dynamic mitochondrial genome of the angiosperm Silene latifolia. BMC Evol Biol 10, 274.

Sloan, D.B., Muller, K., McCauley, D.E., Taylor, D.R., and Storchova, H. (2012a). Intraspecific variation in mitochondrial genome sequence, structure, and gene content in Silene vulgaris, an angiosperm with pervasive cytoplasmic male sterility. New Phytol 196, 1228–1239.

Sloan, D.B., Alverson, A.J., Chuckalovcak, J.P., Wu, M., McCauley, D.E., Palmer, J.D., and Taylor, D.R. (2012b). Rapid Evolution of Enormous, Multichromosomal Genomes in Flowering Plant Mitochondria with Exceptionally High Mutation Rates. Plos Biol 10.

Small, I., Peeters, N., Legeai, F., and Lurin, C. (2004). Predotar: A tool for rapidly screening proteomes for N-terminal targeting sequences. Proteomics 4, 1581–1590.

Small, I., Marechal-Drouard, L., Masson, J., Pelletier, G., Cosset, A., Weil, J.H., and Dietrich, A. (1992). In vivo import of a normal or mutagenized heterologous transfer RNA into the mitochondria of transgenic plants: towards novel ways of influencing mitochondrial gene expression? EMBO J 11, 1291–1296.

Small, I., Akashi, K., Chapron, A., Dietrich, A., Duchêne, A.M., Lancelin, D., Marechal-Drouard, L., Menand, B., Mireau, H., Moudden, Y., Ovesna, J., Peeters, N., Sakamoto, W., Souciet, G., and Wintz, H. (1999). The strange evolutionary history of plant mitochondrial tRNAs and their aminoacyl-tRNA synthetases. J Hered 90, 333–337.

Sperschneider, J., Catanzariti, A.M., DeBoer, K., Petre, B., Gardiner, D.M., Singh, K.B., Dodds, P.N., and Taylor, J.M. (2017). LOCALIZER: subcellular localization prediction of both plant and effector proteins in the plant cell. Sci Rep 7, 44598.

Stamatakis, A. (2014). RAxML version 8: a tool for phylogenetic analysis and post-analysis of large phylogenies. Bioinformatics 30, 1312–1313.

Timmis, J.N., Ayliffe, M.A., Huang, C.Y., and Martin, W. (2004). Endosymbiotic gene transfer: organelle genomes forge eukaryotic chromosomes. Nat Rev Genet 5, 123–135.

Tsunoda, M., Kusakabe, Y., Tanaka, N., Ohno, S., Nakamura, M., Senda, T., Moriguchi, T., Asai, N., Sekine, M., and Yokogawa, T. (2007). Structural basis for recognition of cognate tRNA by tyrosyl-tRNA synthetase from three kingdoms. Nucleic Acids Res 35, 4289–4300.

Warren, J.M., and Sloan, D.B. (2020). Interchangeable parts: The evolutionarily dynamic tRNA population in plant mitochondria. Mitochondrion 52, 144–156.

Warren, J.M., Salinas-Giegé, T., Triant, D.A., Taylor, D.R., Drouard, L., and Sloan, D.B. (2021). Rapid shifts in mitochondrial tRNA import in a plant lineage with extensive mitochondrial tRNA gene loss. Mol Biol Evol.

Williams, A.M., Itgen, M.W., Broz, A.K., Carter, O.G., and Sloan, D.B. (2021). Long-read transcriptome and other genomic resources for the angiosperm Silene noctiflora. G3 (Bethesda) 11.

Williams, E.J., Pal, C., and Hurst, L.D. (2000). The molecular evolution of signal peptides. Gene 253, 313–322.

Wu, Z., Sloan, D.B., Brown, C.W., Rosenblueth, M., Palmer, J.D., and Ong, H.C. (2017). Mitochondrial Retroprocessing Promoted Functional Transfers of rpl5 to the Nucleus in Grasses. Mol Biol Evol 34, 2340–2354.

Yarham, J.W., Elson, J.L., Blakely, E.L., McFarland, R., and Taylor, R.W. (2010). Mitochondrial tRNA mutations and disease. Wiley Interdiscip Rev RNA 1, 304–324.

Yu, G. (2020). Using ggtree to Visualize Data on Tree-Like Structures. Curr Protoc Bioinformatics 69, e96.

Zhao, L., Zhang, H., Kohnen, M.V., Prasad, K., Gu, L., and Reddy, A.S.N. (2019). Analysis of Transcriptome and Epitranscriptome in Plants Using PacBio Iso-Seq and Nanopore-Based Direct RNA Sequencing. Front Genet 10, 253.

