## Supplementary figures and images for "Rewiring of aminoacyl-tRNA synthetase localization and interactions in plants with extensive mitochondrial tRNA gene loss"

### Supplemental Figure 1

##### Cytosolic/Organellar AlaRS

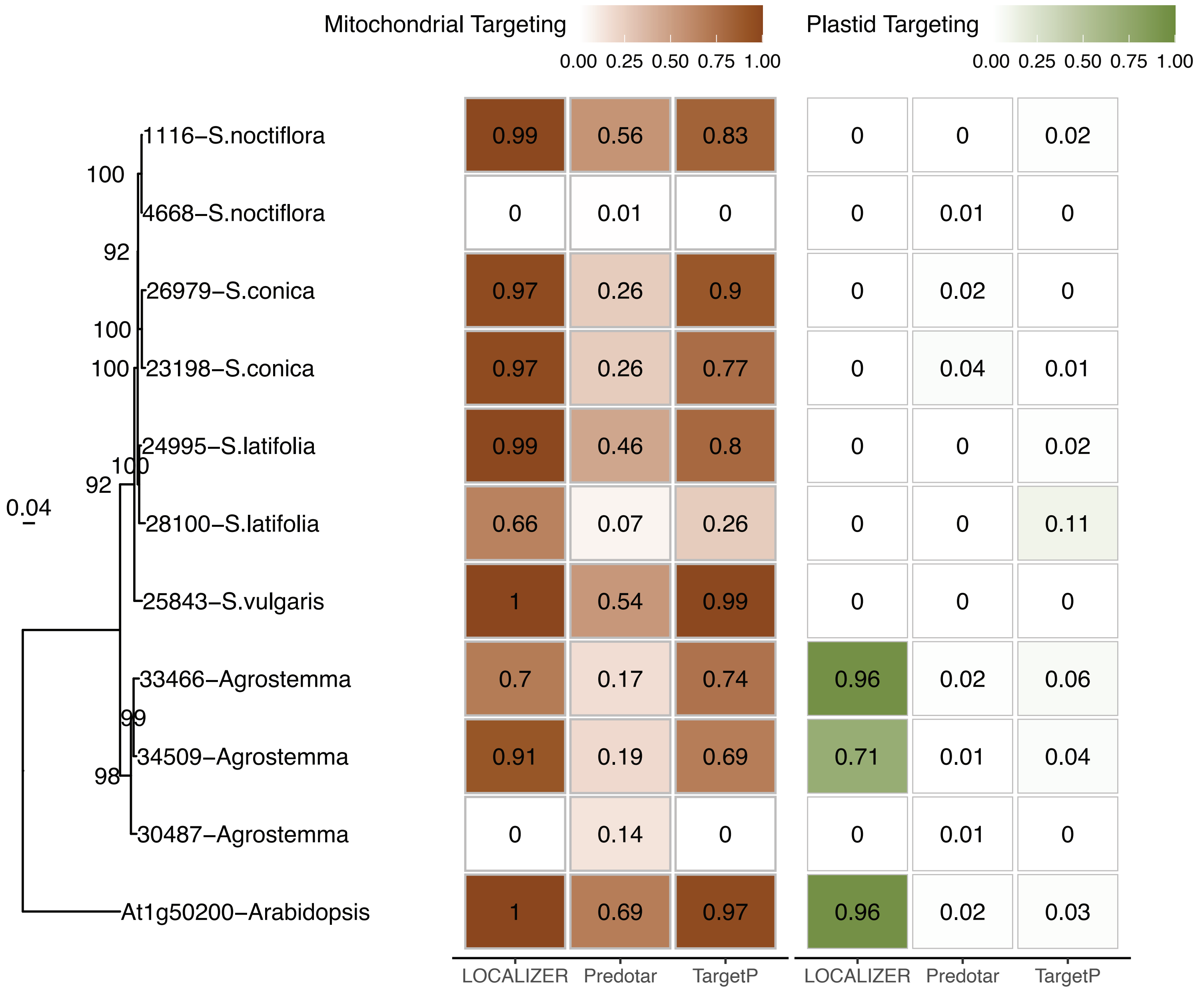


##### Organellar AlaRS

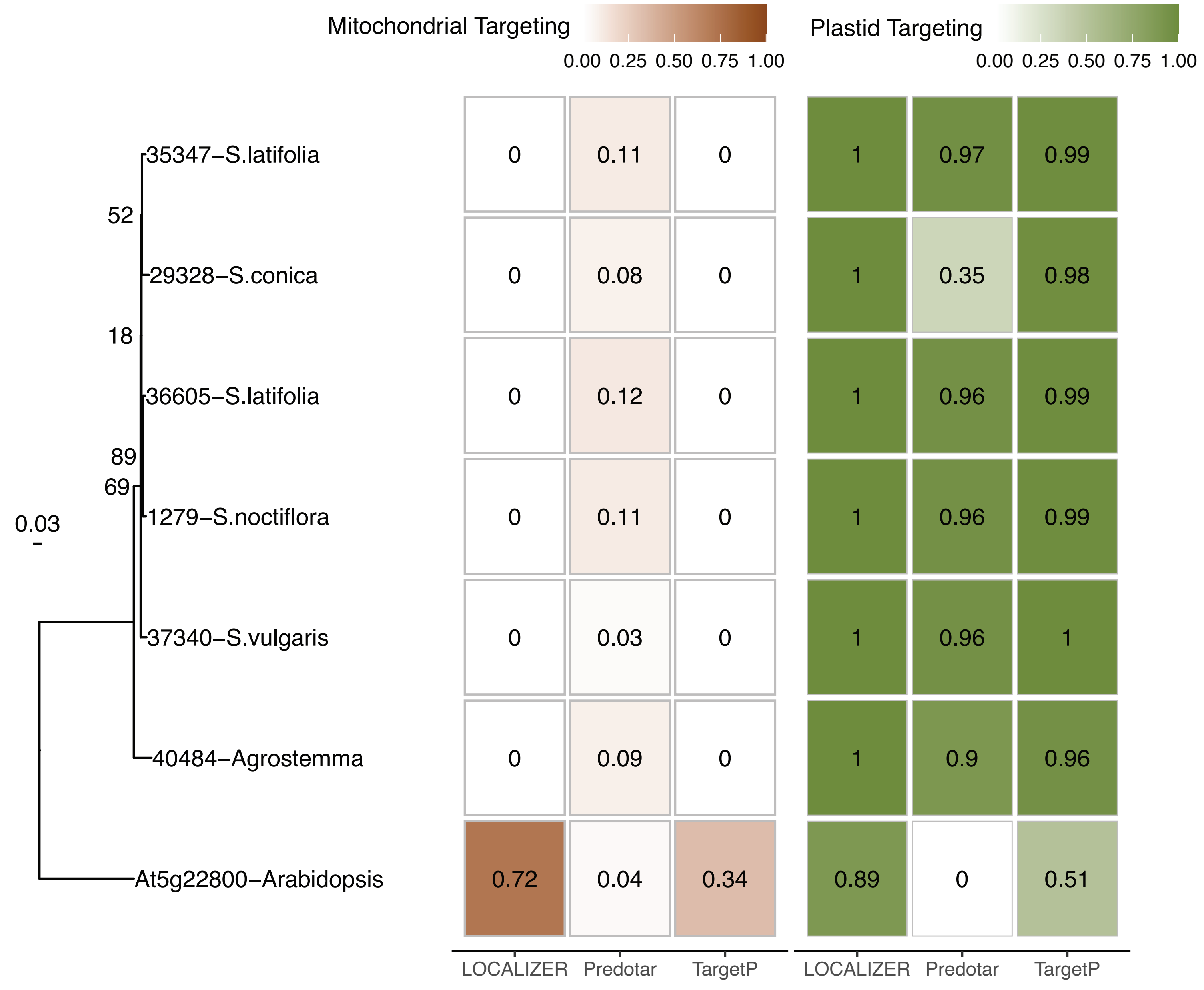

### Supplemental Figure 1

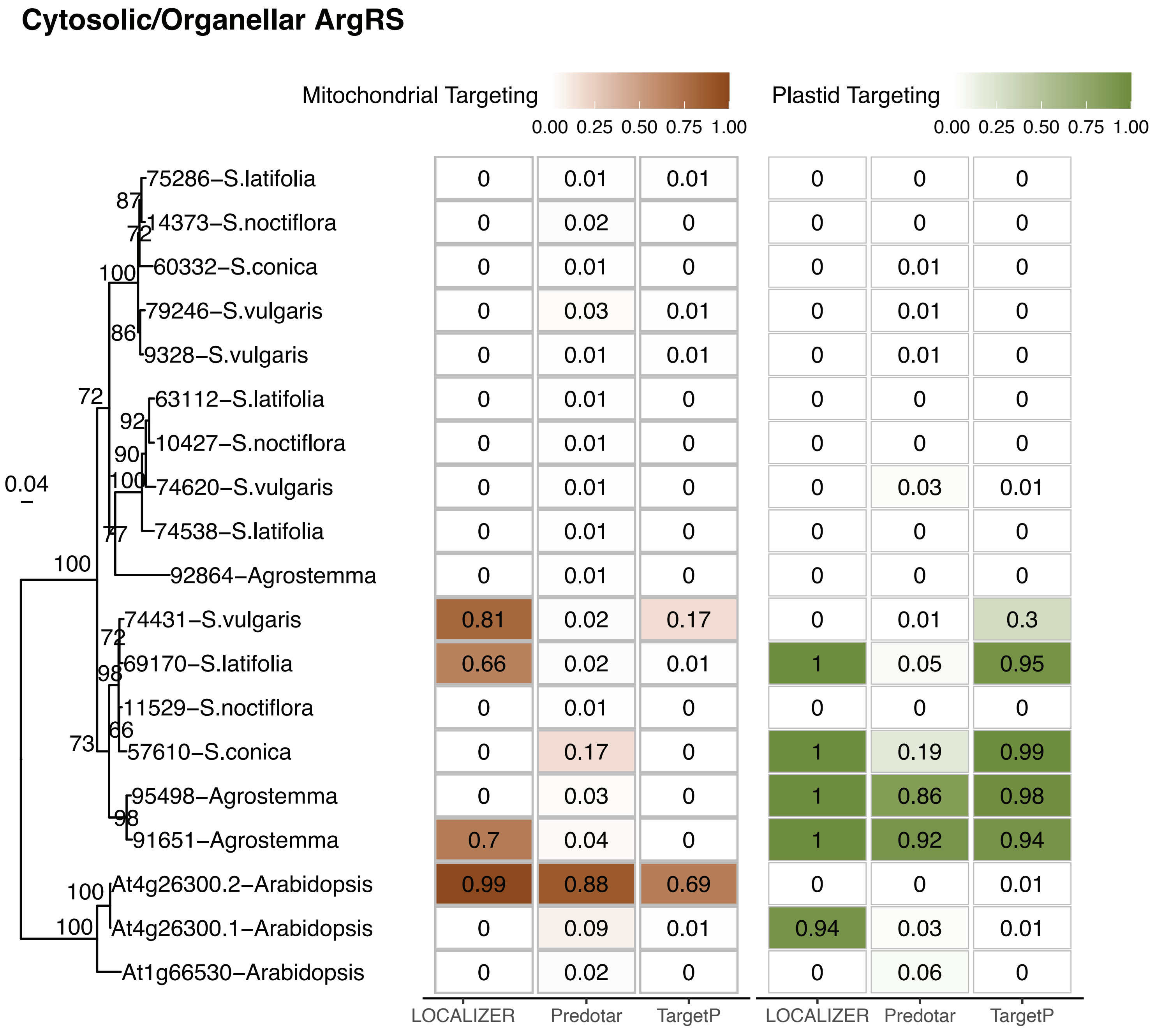

### Supplemental Figure 2

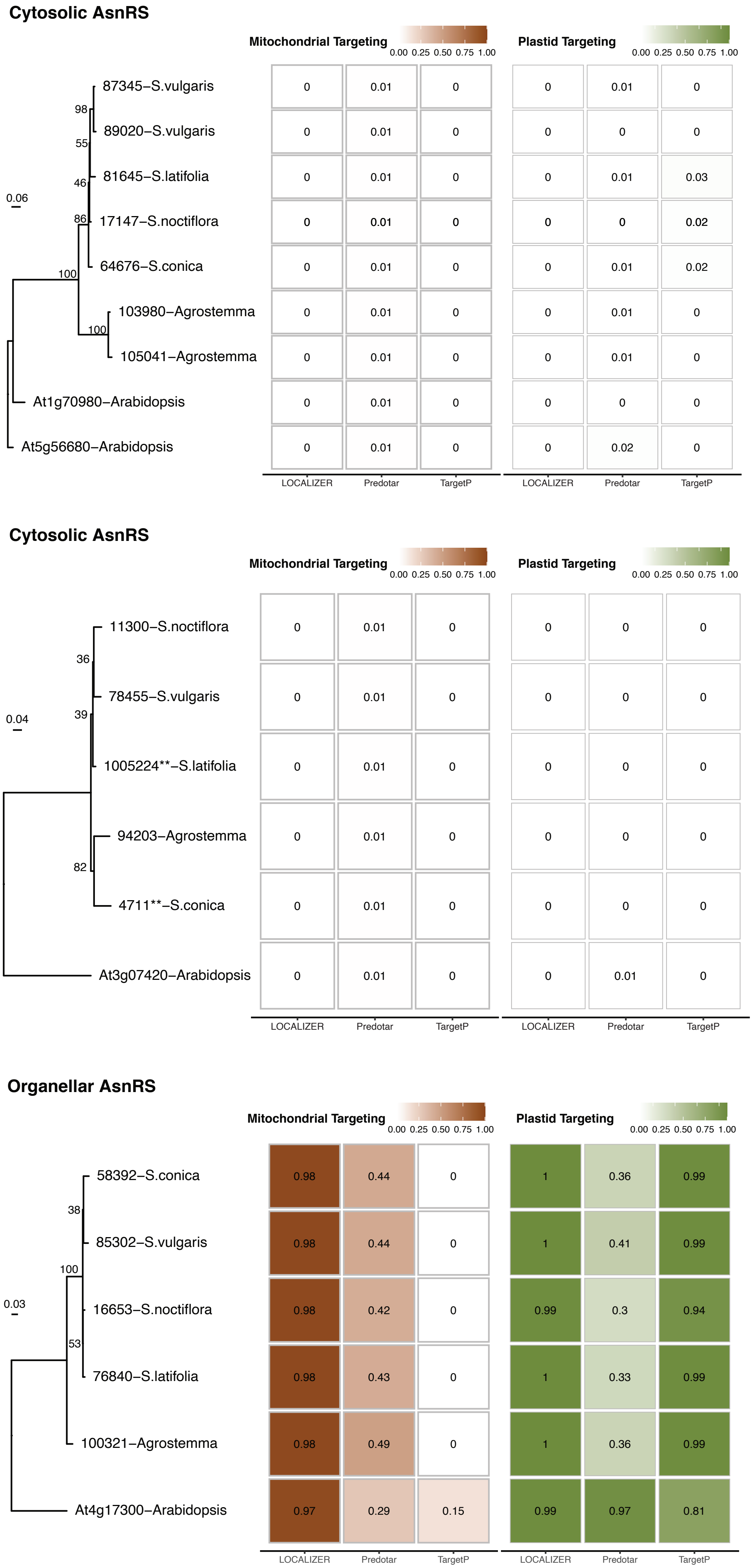

### Supplemental Figure 3

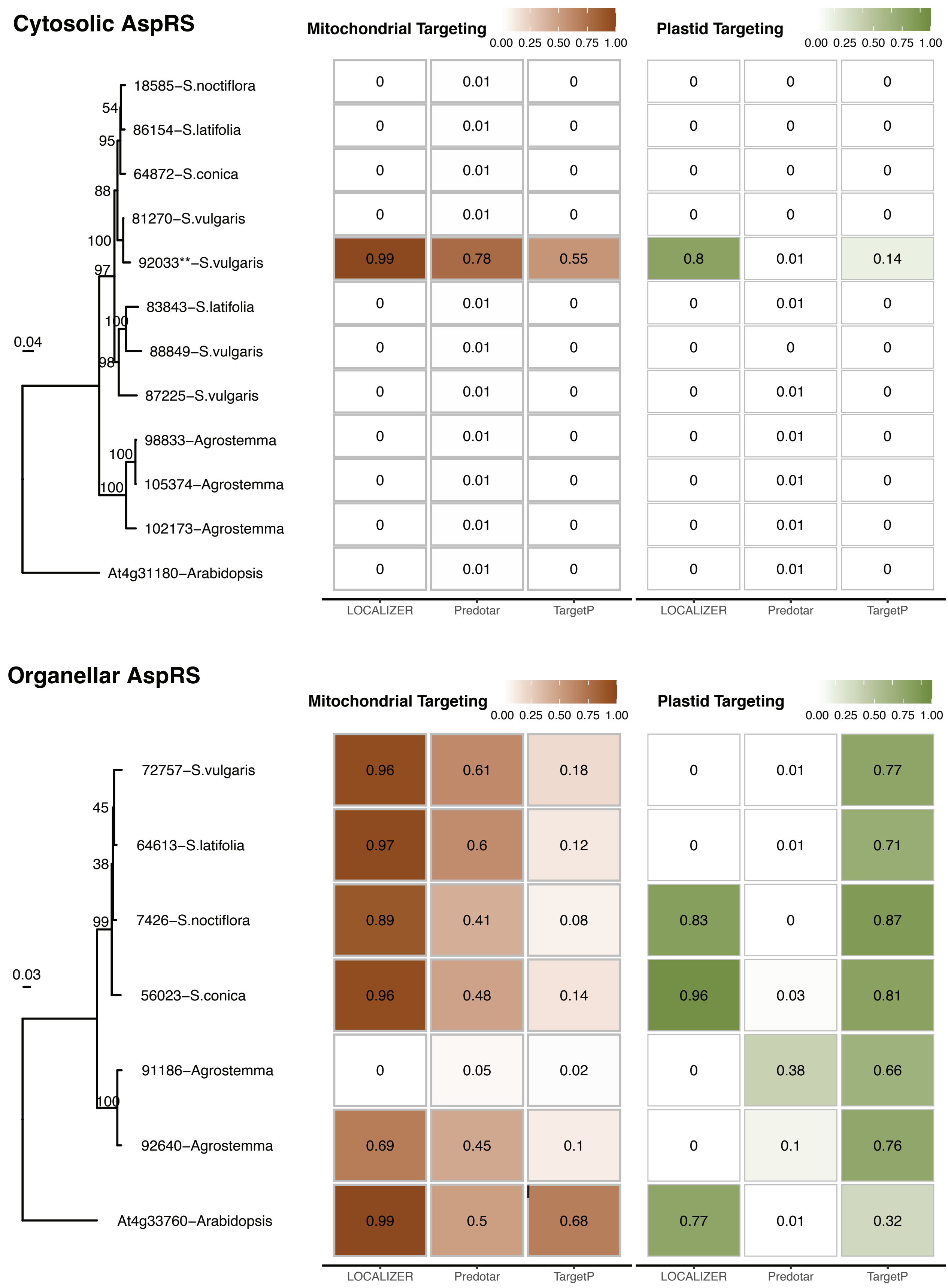

### Supplemental Figure 4

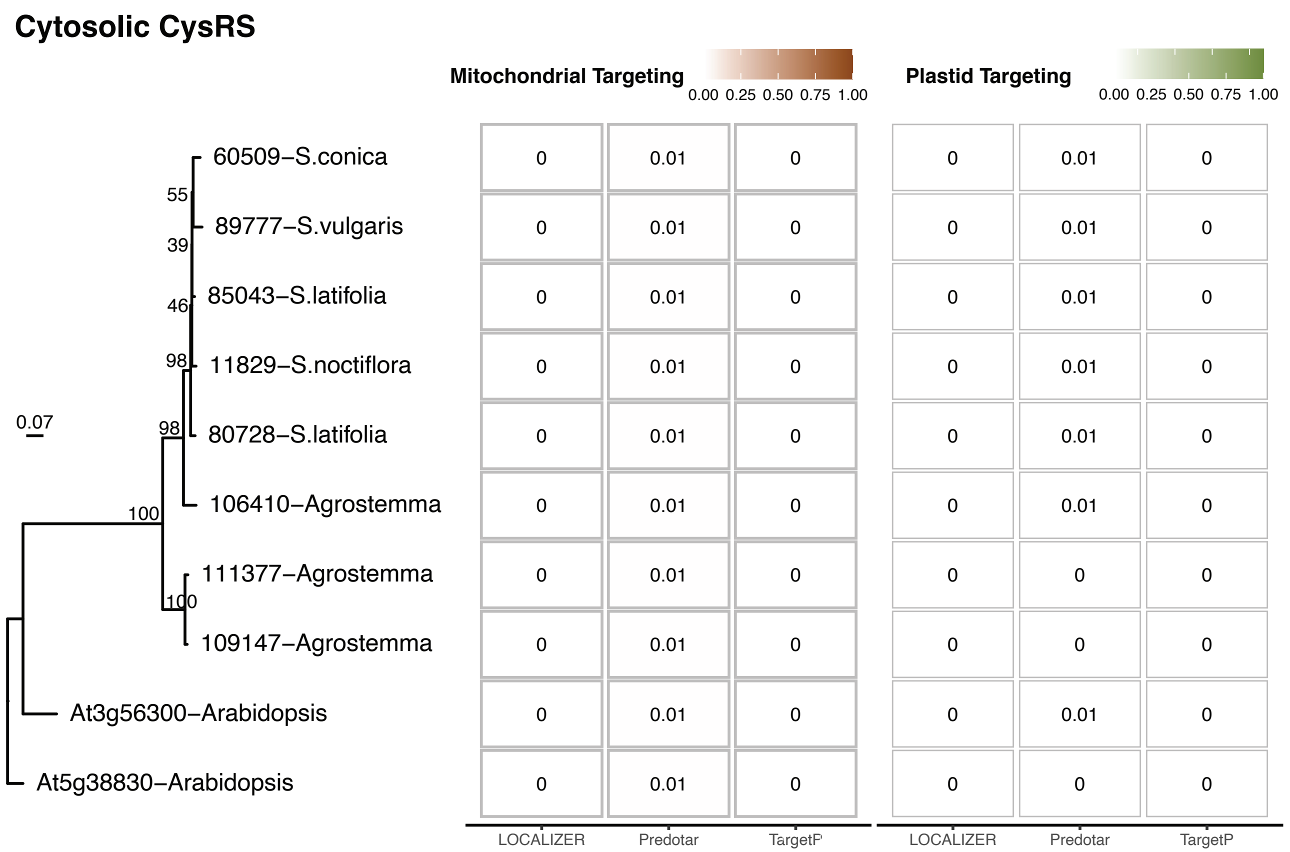

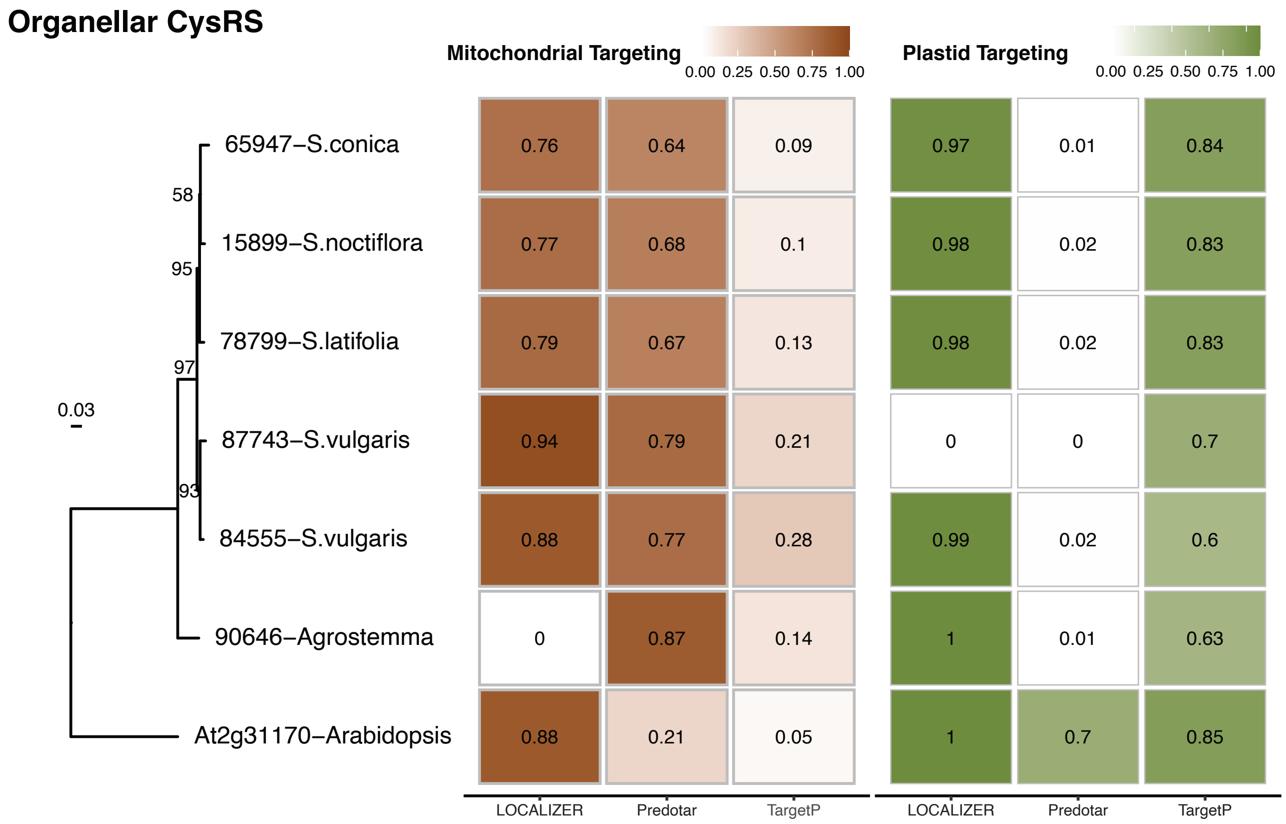

### Supplemental Figure 5

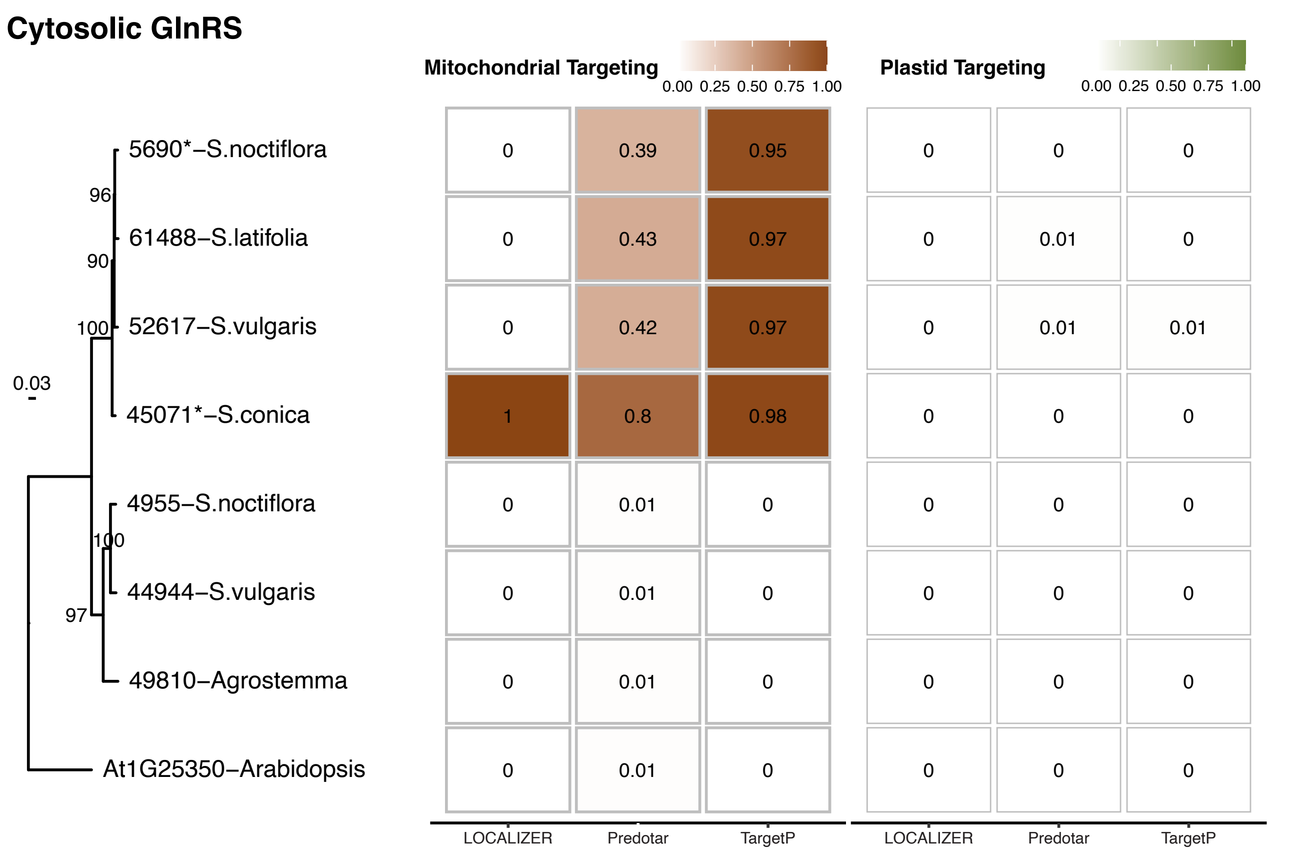

### Supplemental Figure 6

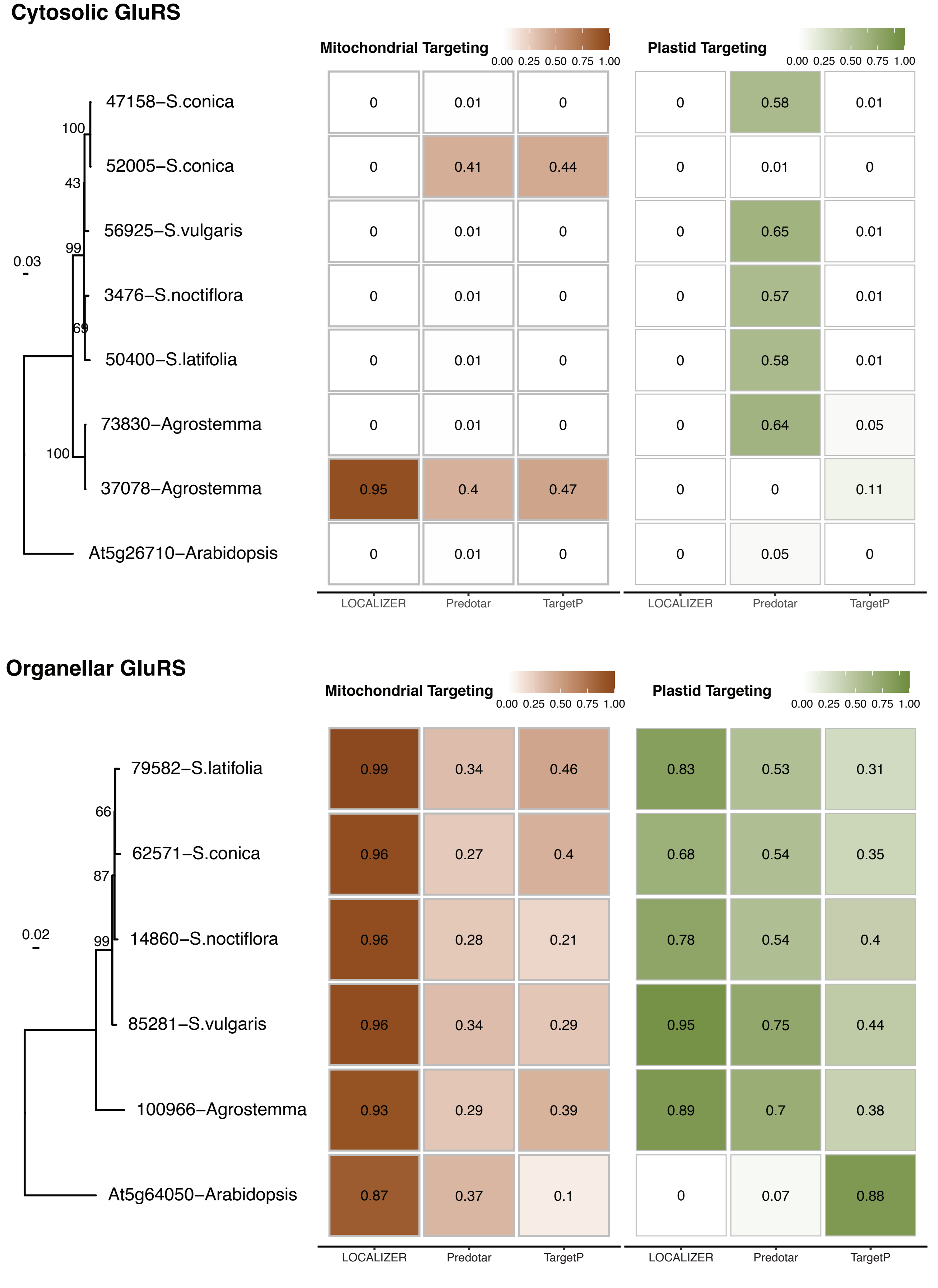

### Supplemental Figure 7

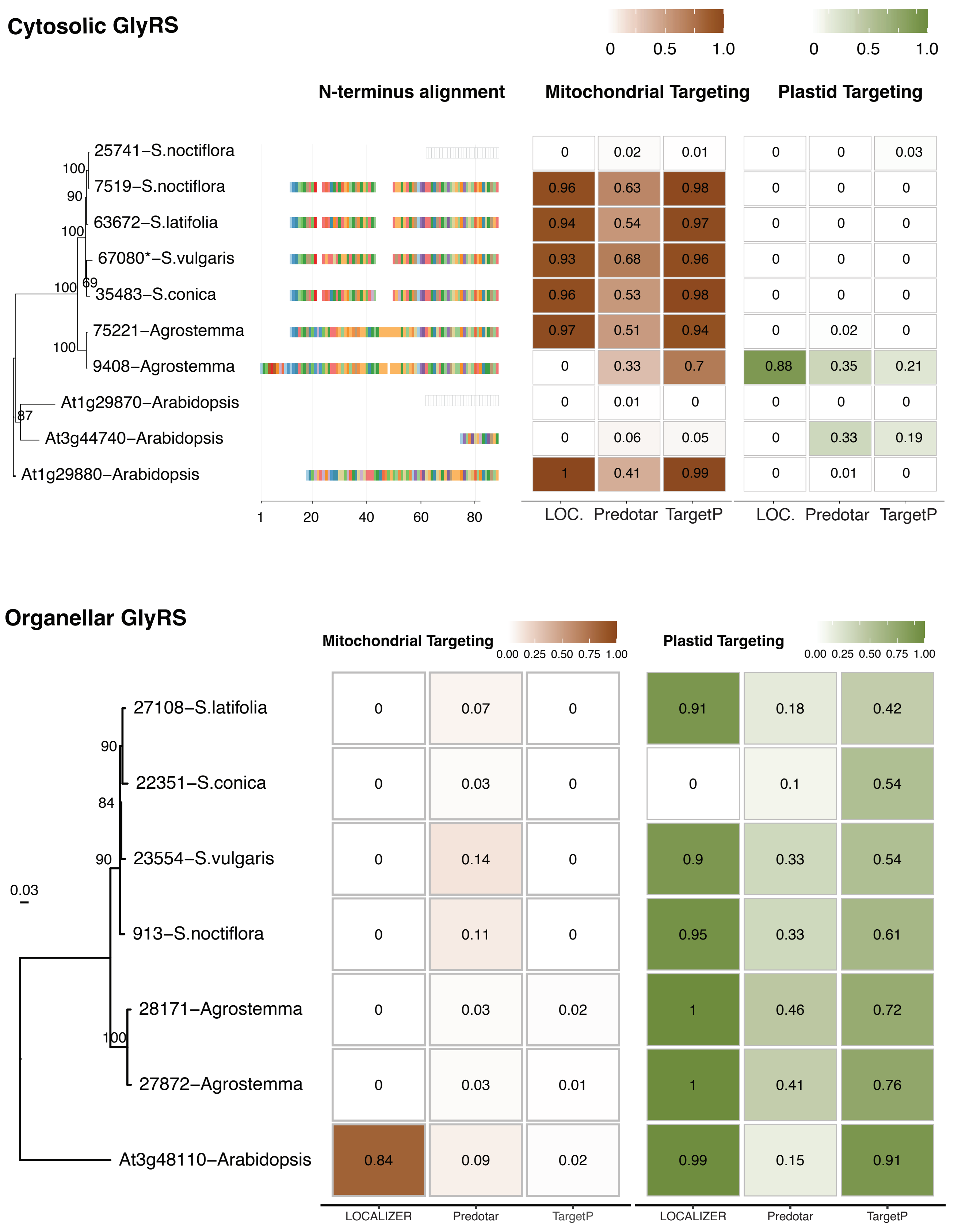

### Supplemental Figure 8

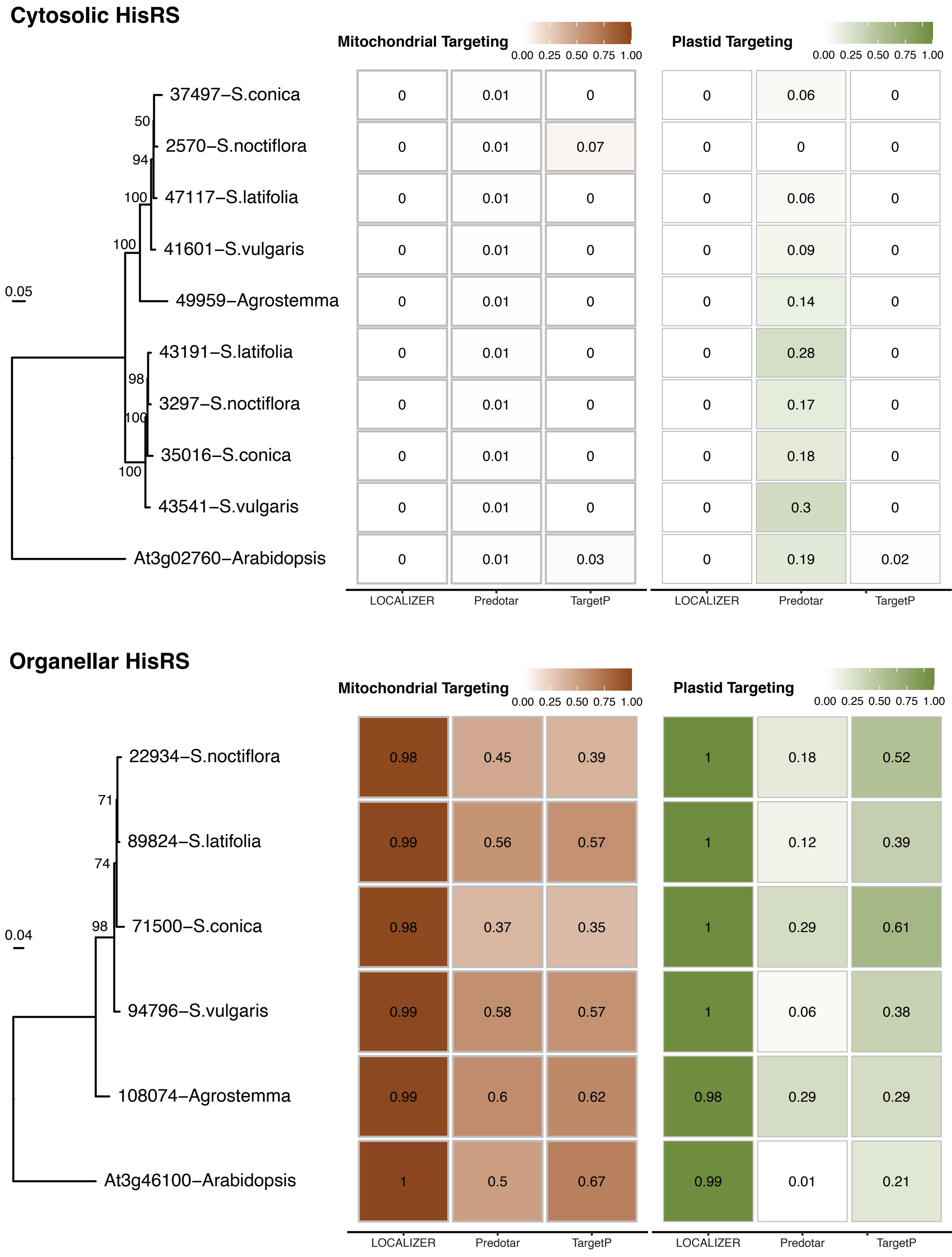

### Supplemental Figure 9

##### Cytosolic IleRS

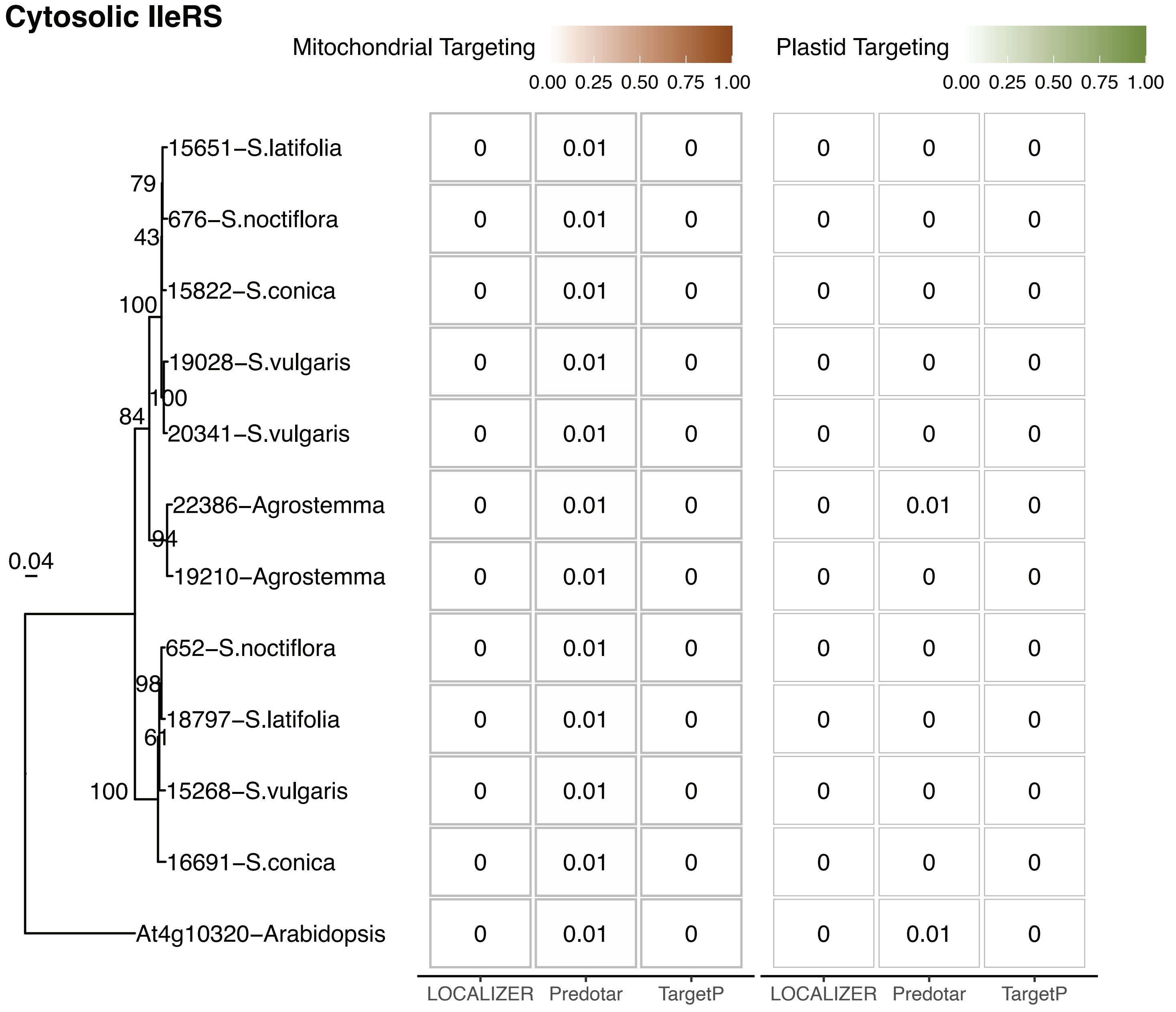


##### Organellar IleRS

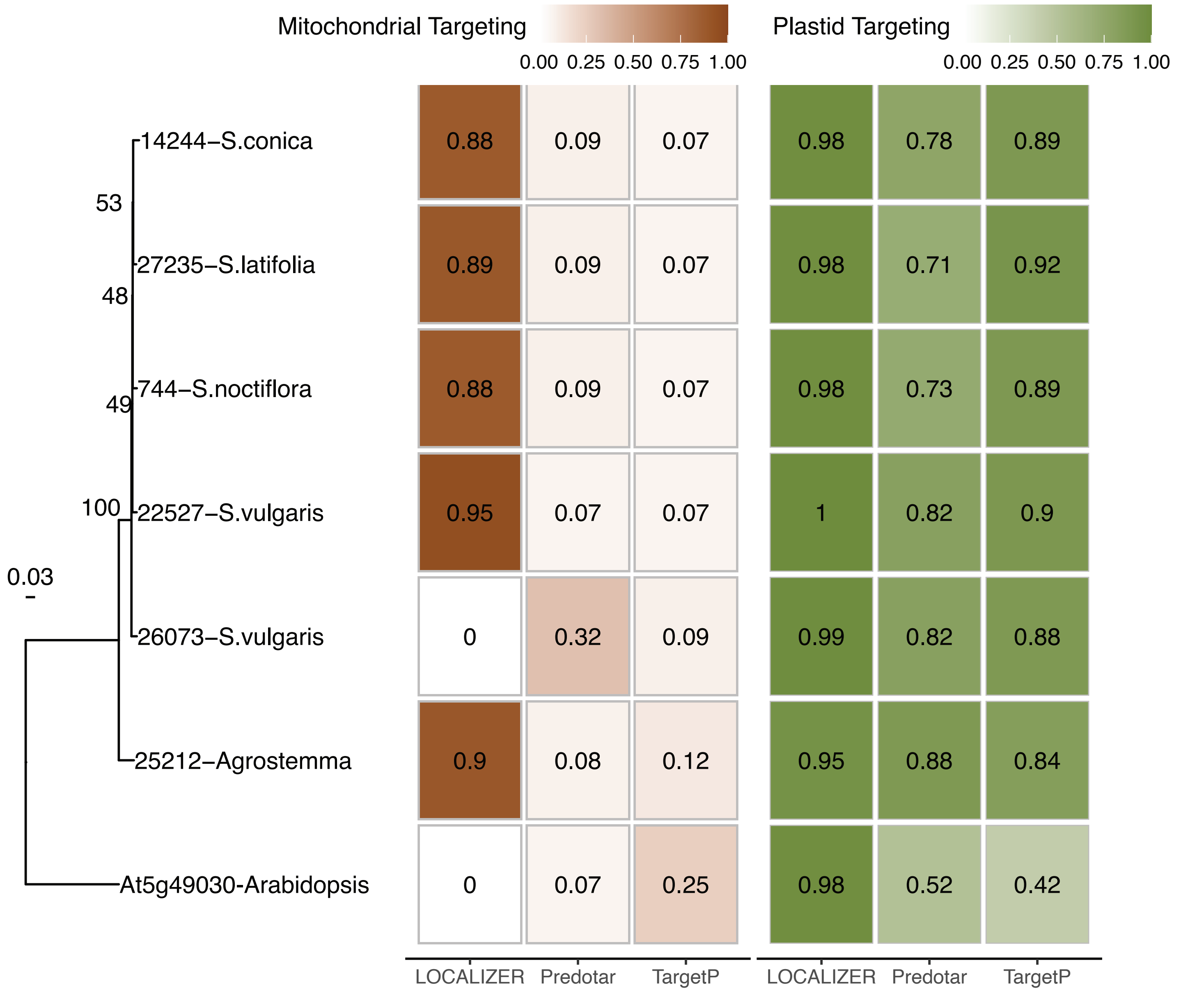

### Supplemental Figure 11

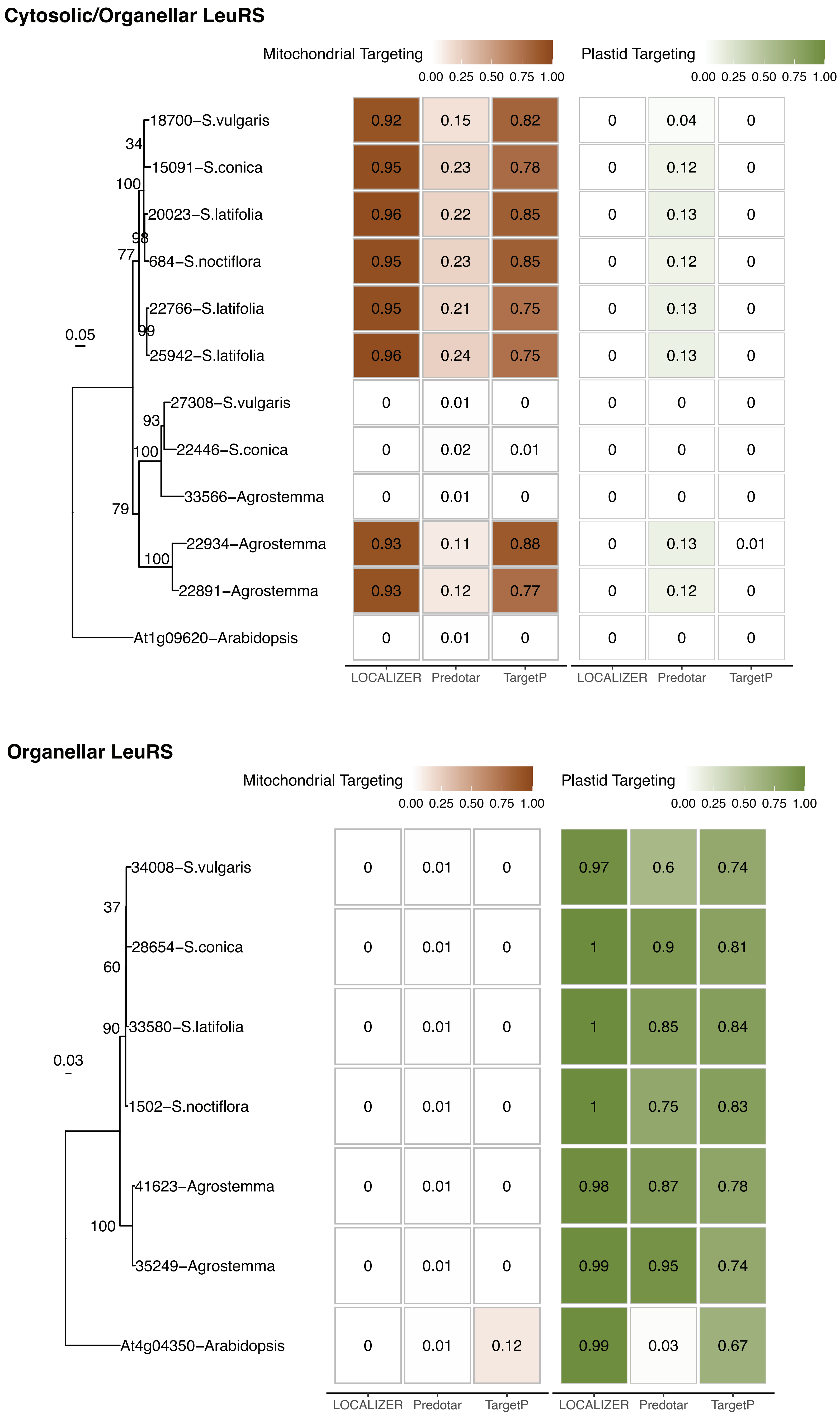

### Supplemental Figure 12

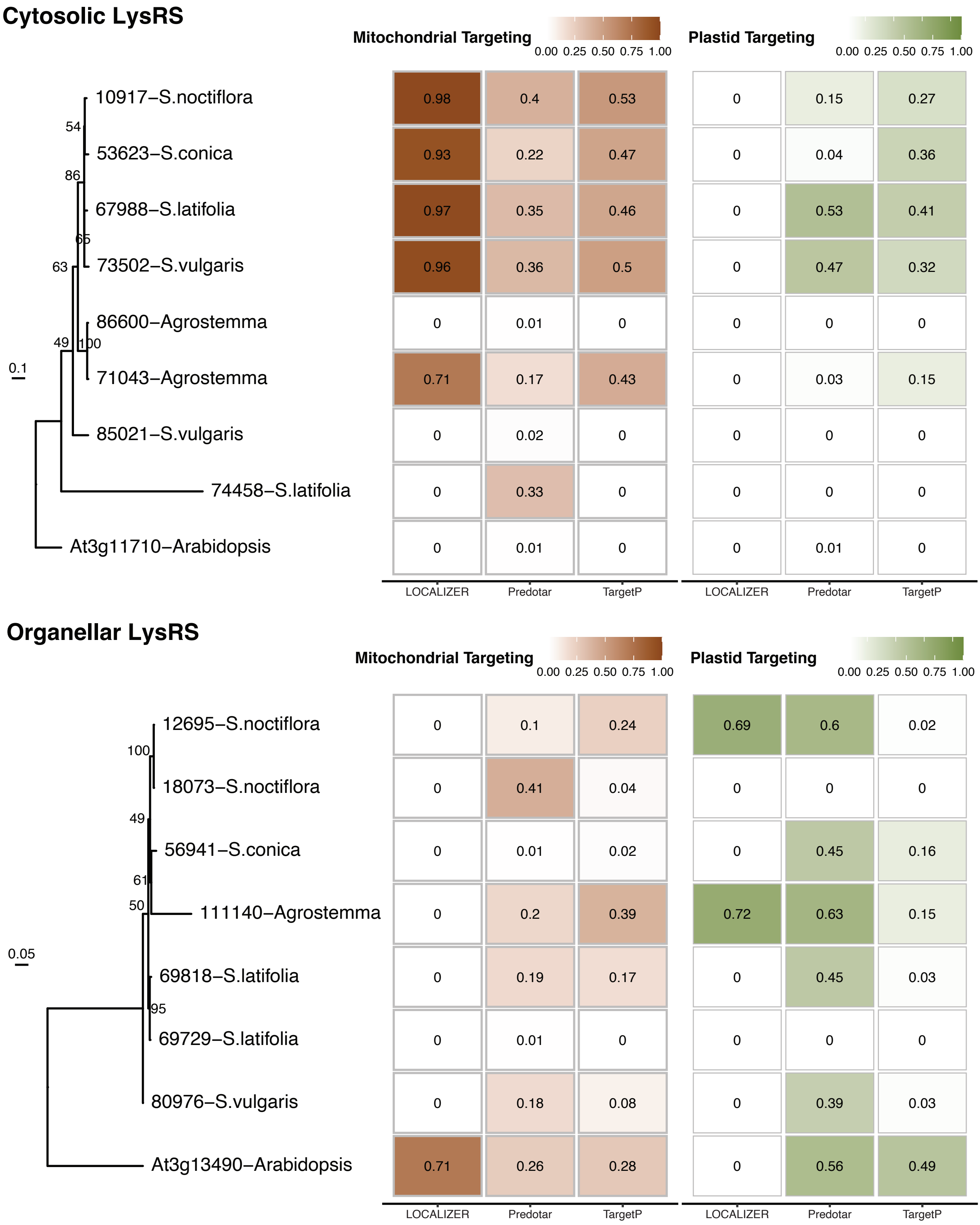

### Supplemental Figure 13

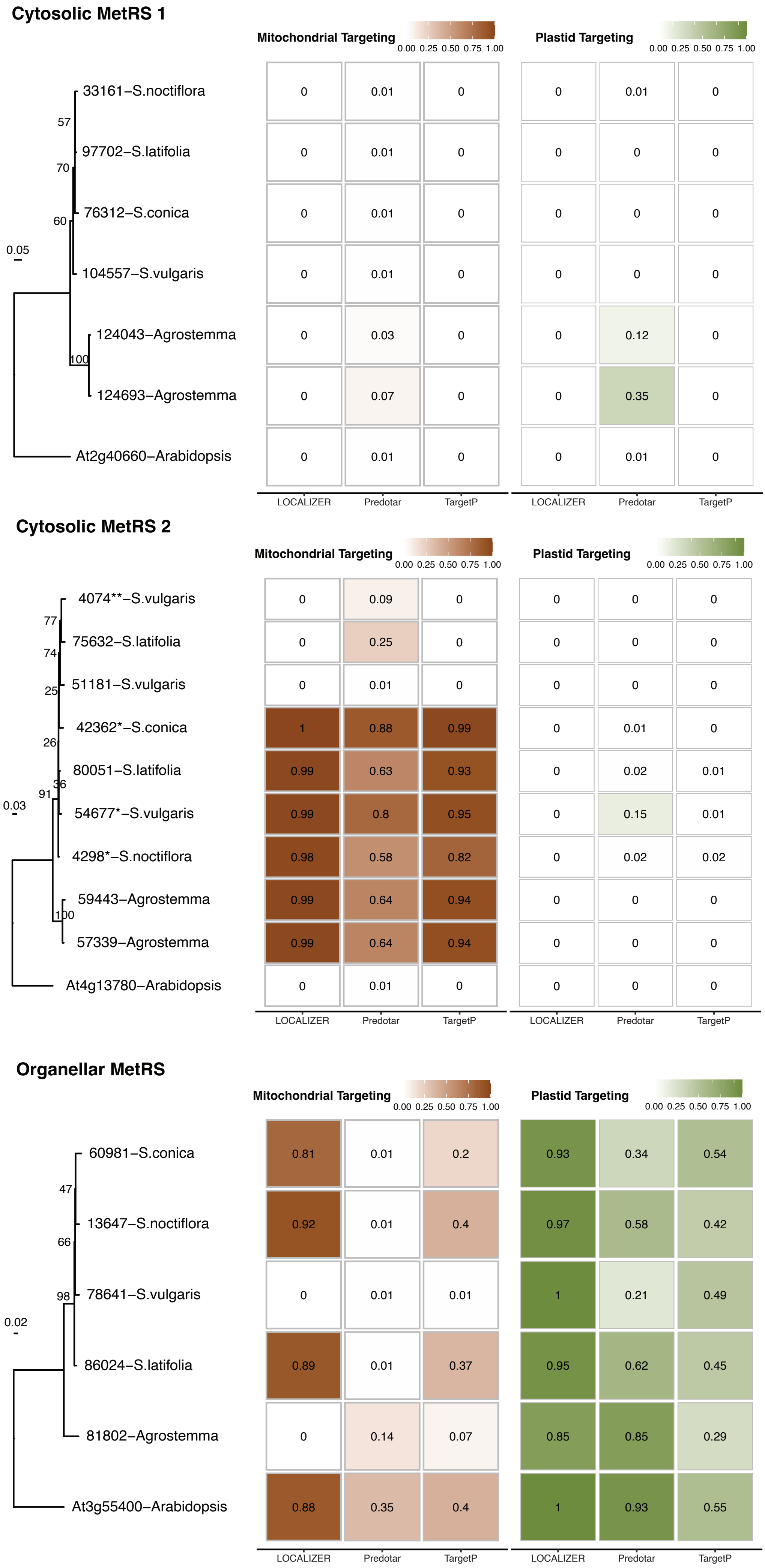

### Supplemental Figure 14

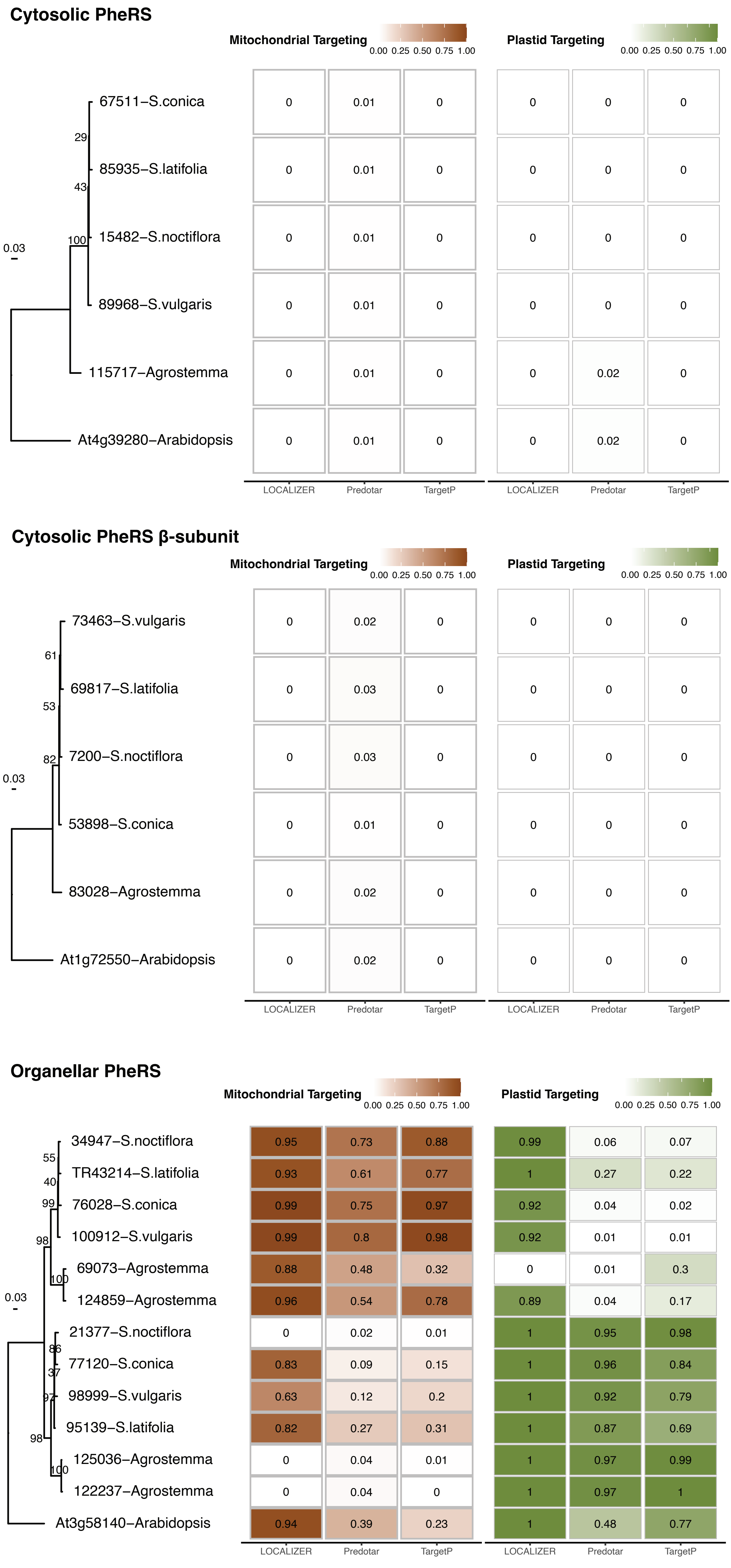

### Supplemental Figure 15

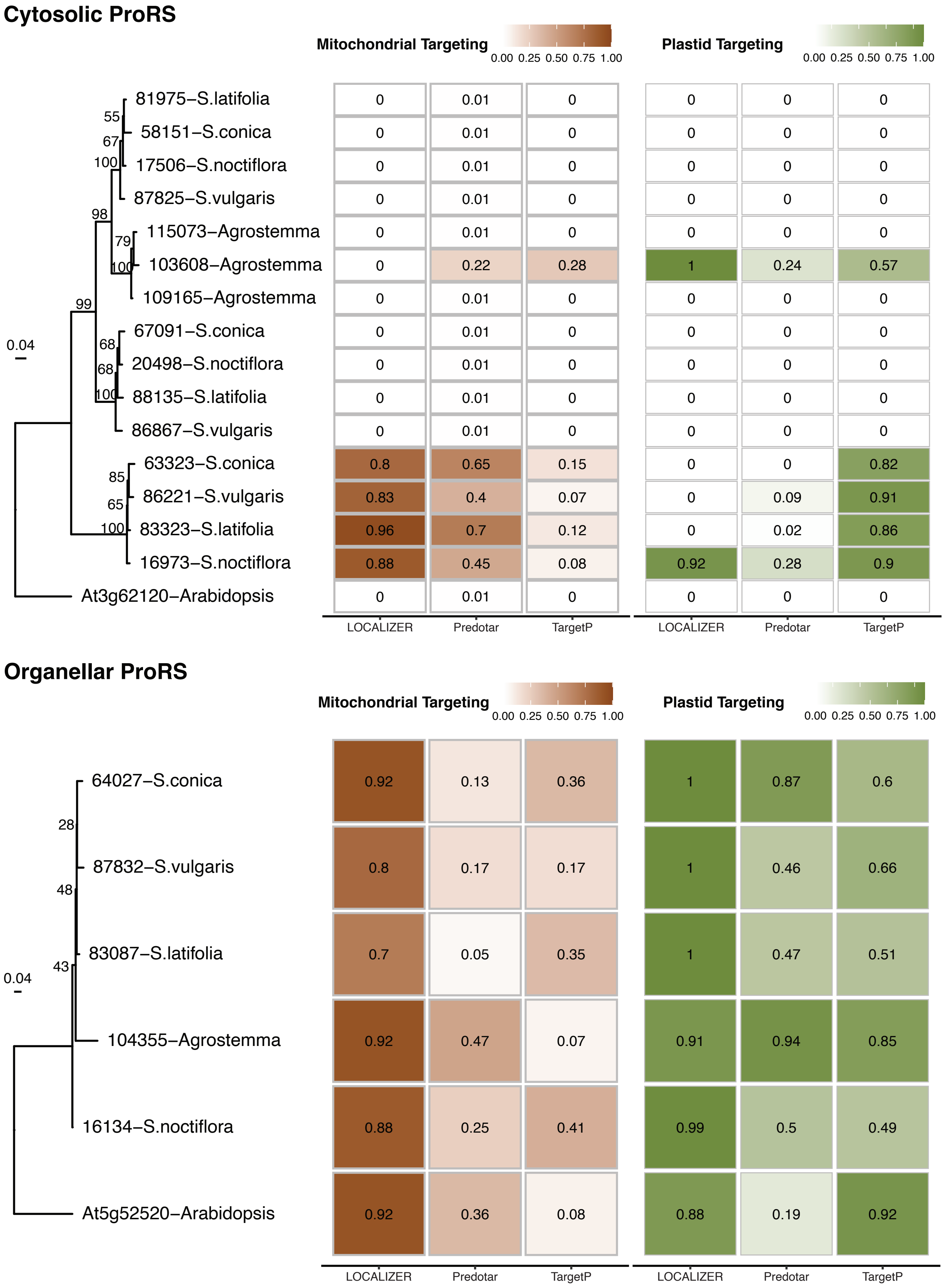

### Supplemental Figure 16

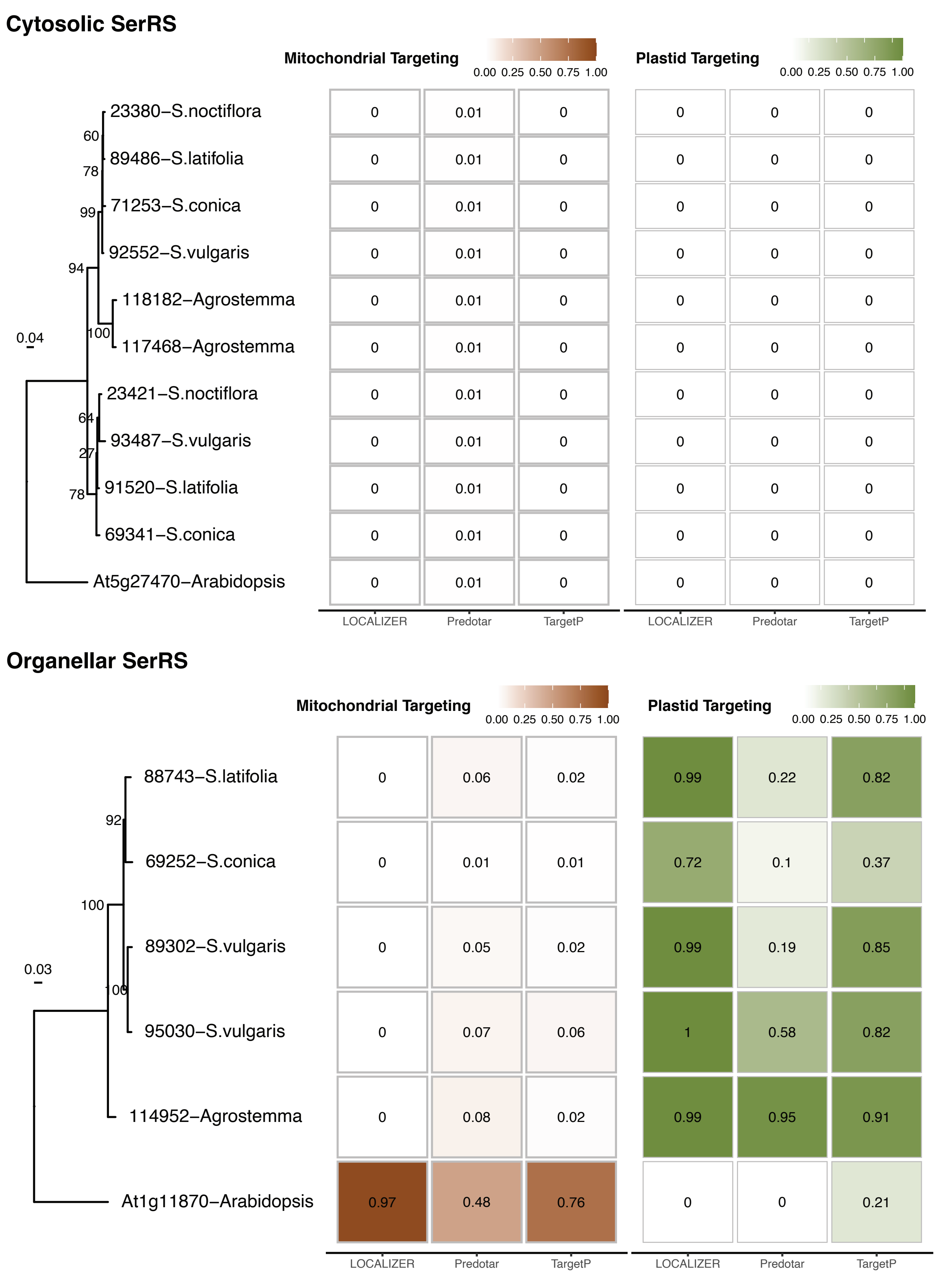

### Supplemental Figure 17

##### Cytosolic/Organellar ThrRS

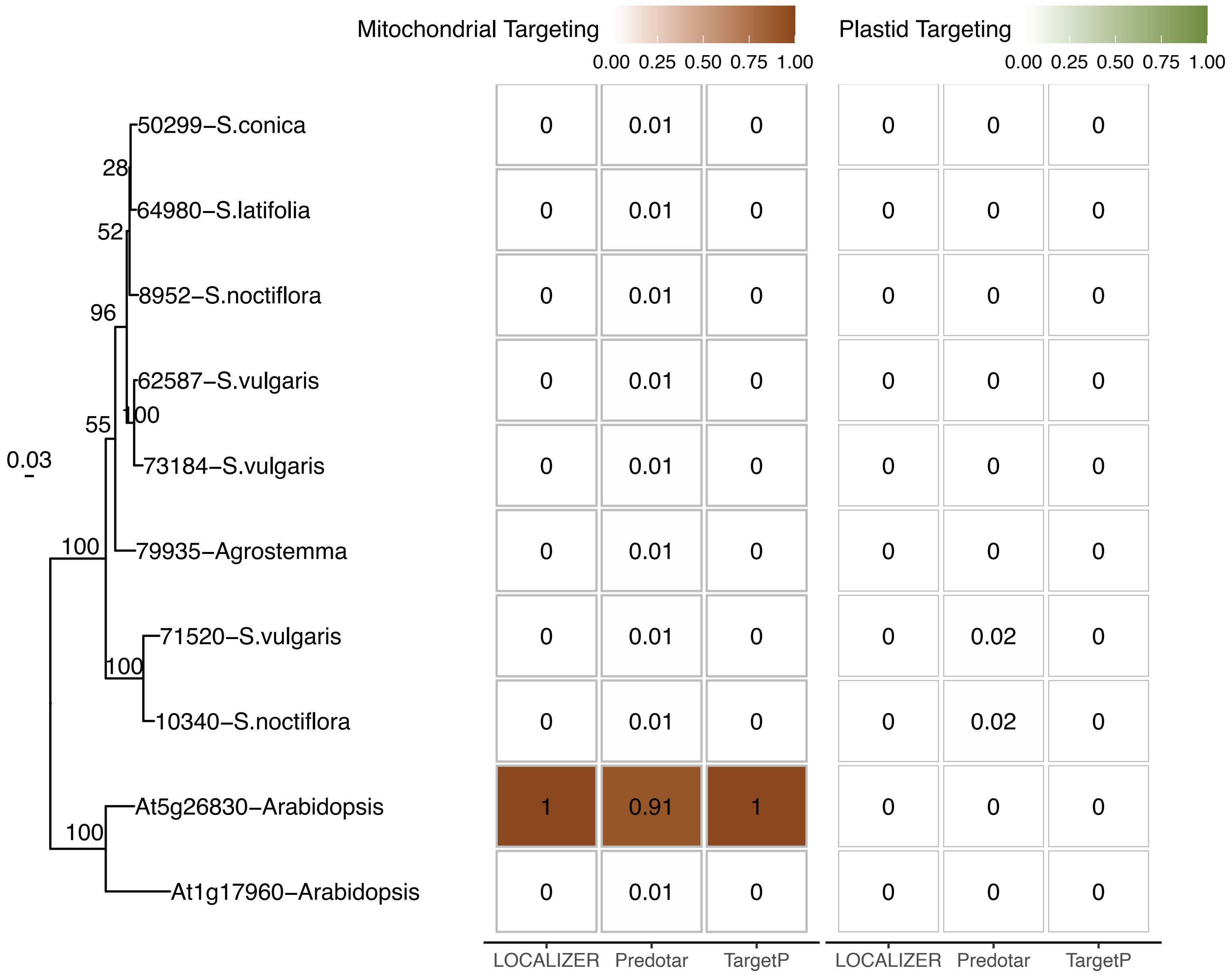


##### Organellar ThrRS

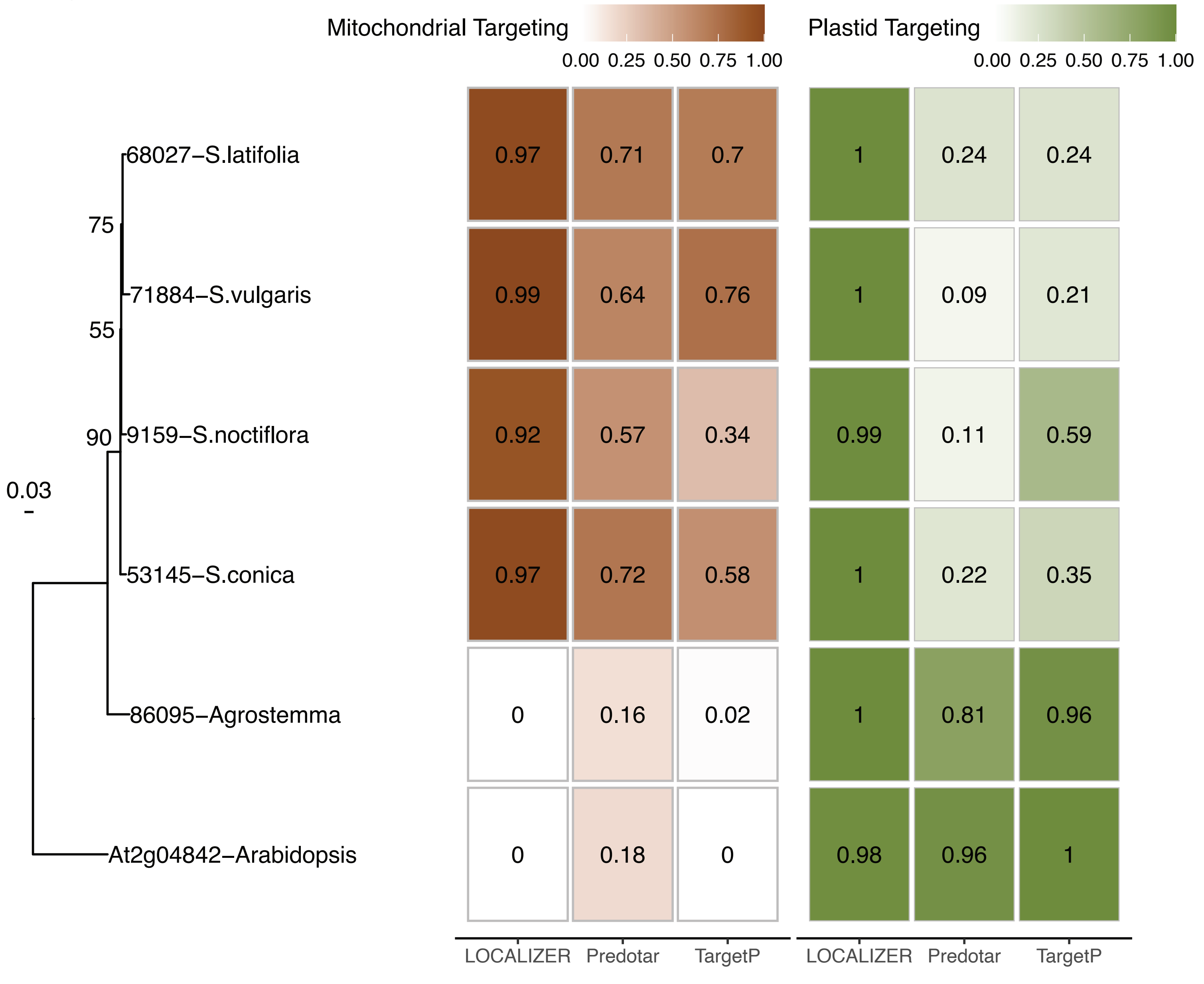

### Supplemental Figure 18

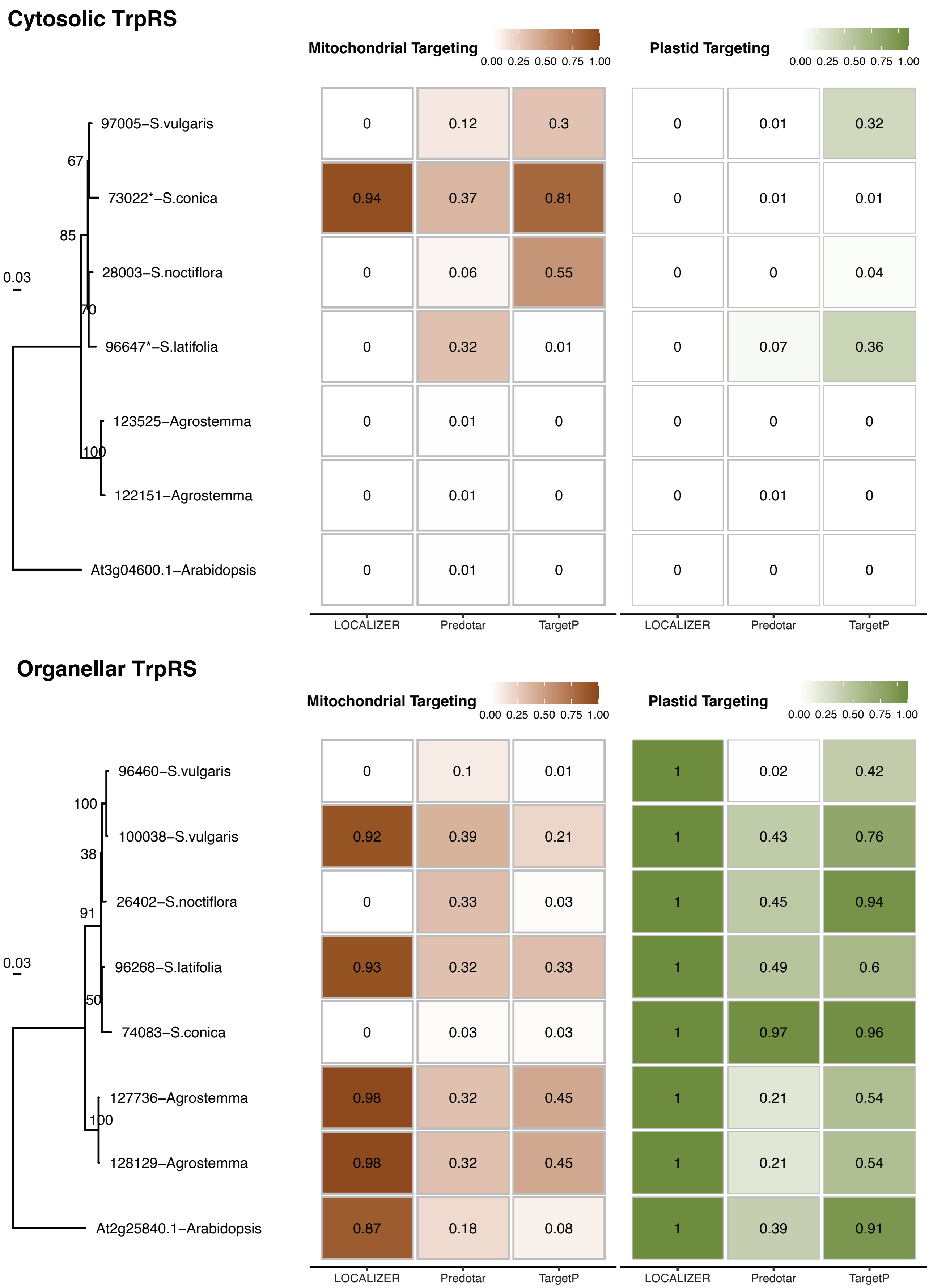

### Supplemental Figure 19

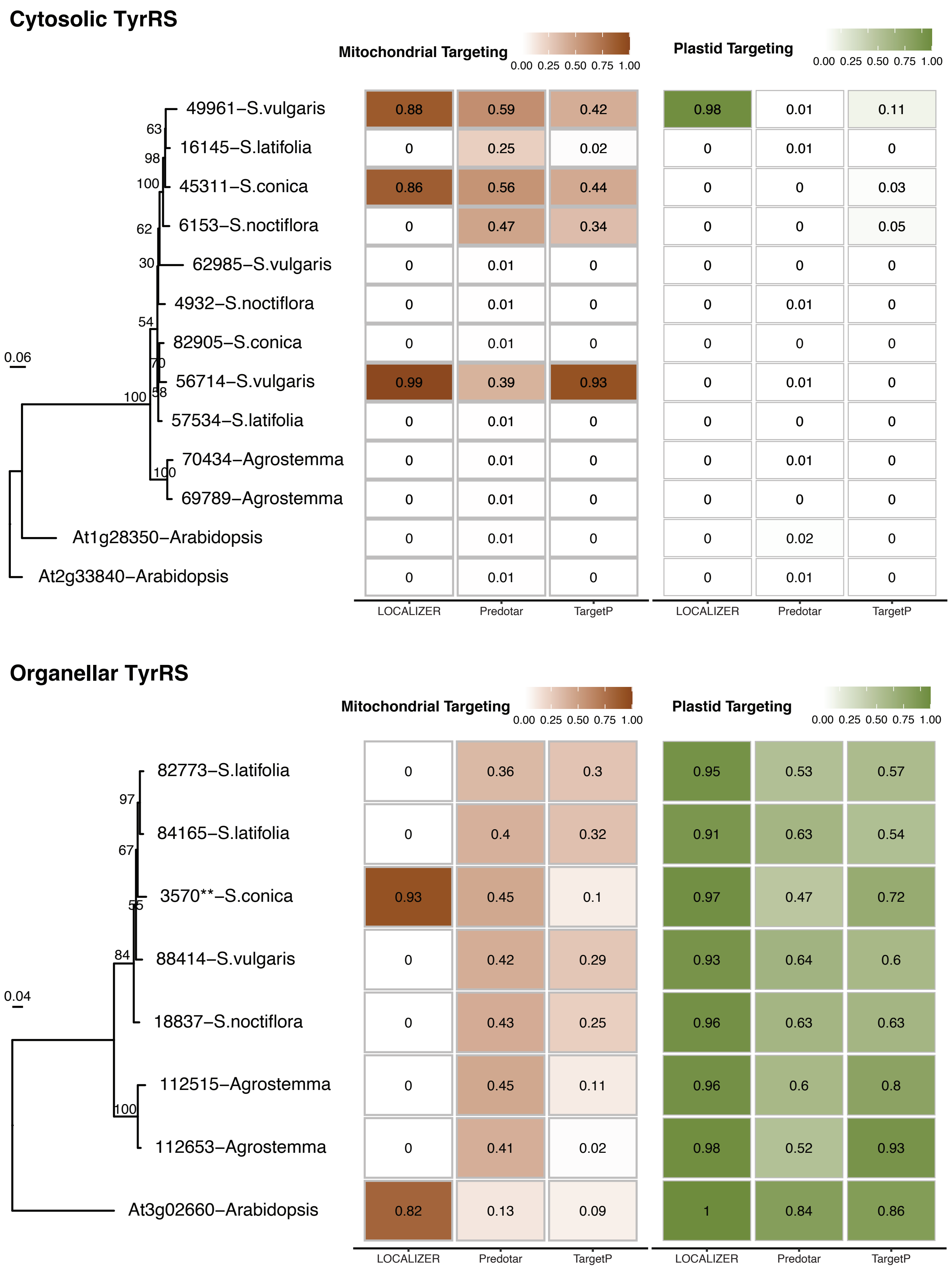

### Supplemental Figure 20

##### Cytosolic/Organellar ValRS

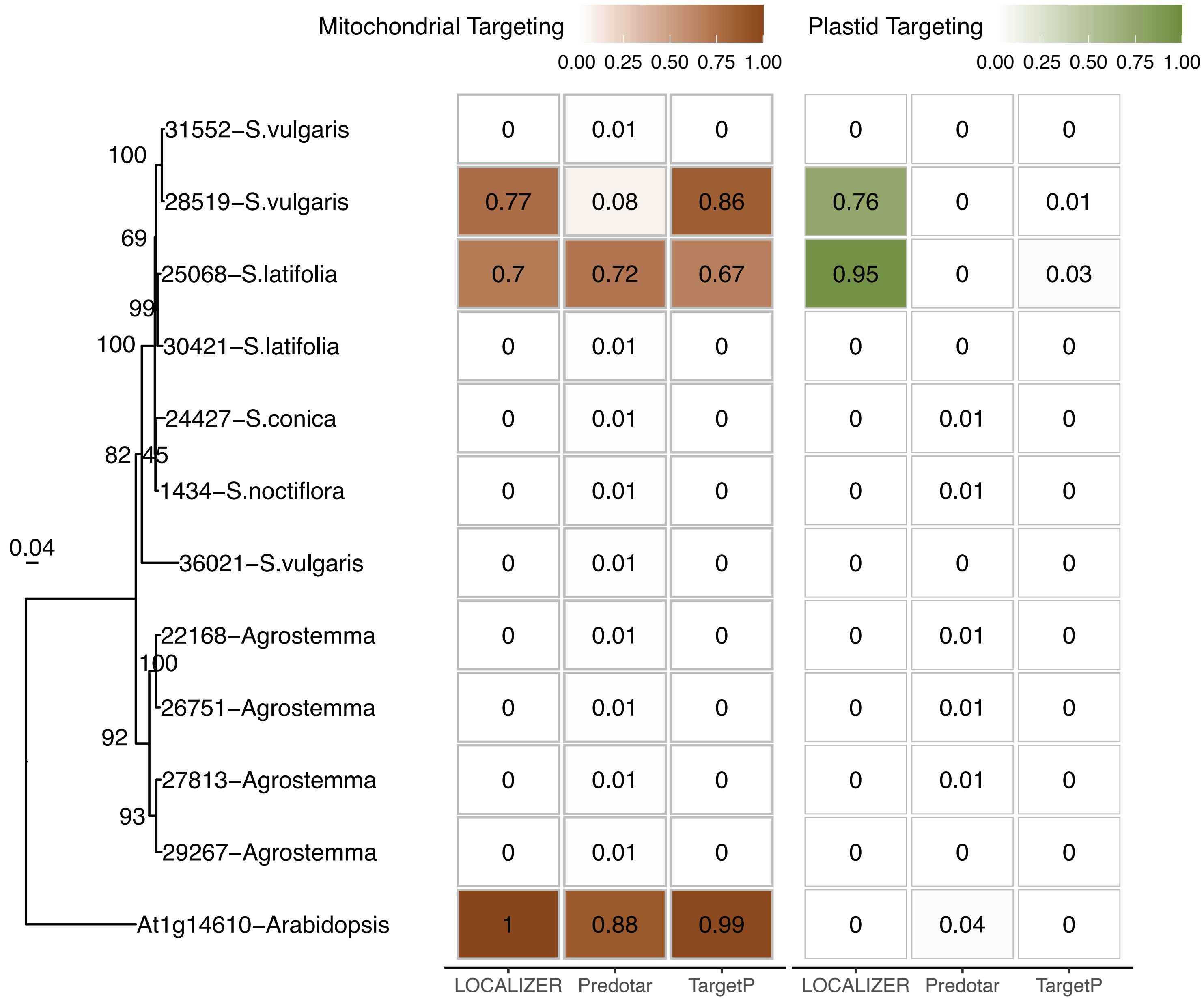


##### Organellar ValRS

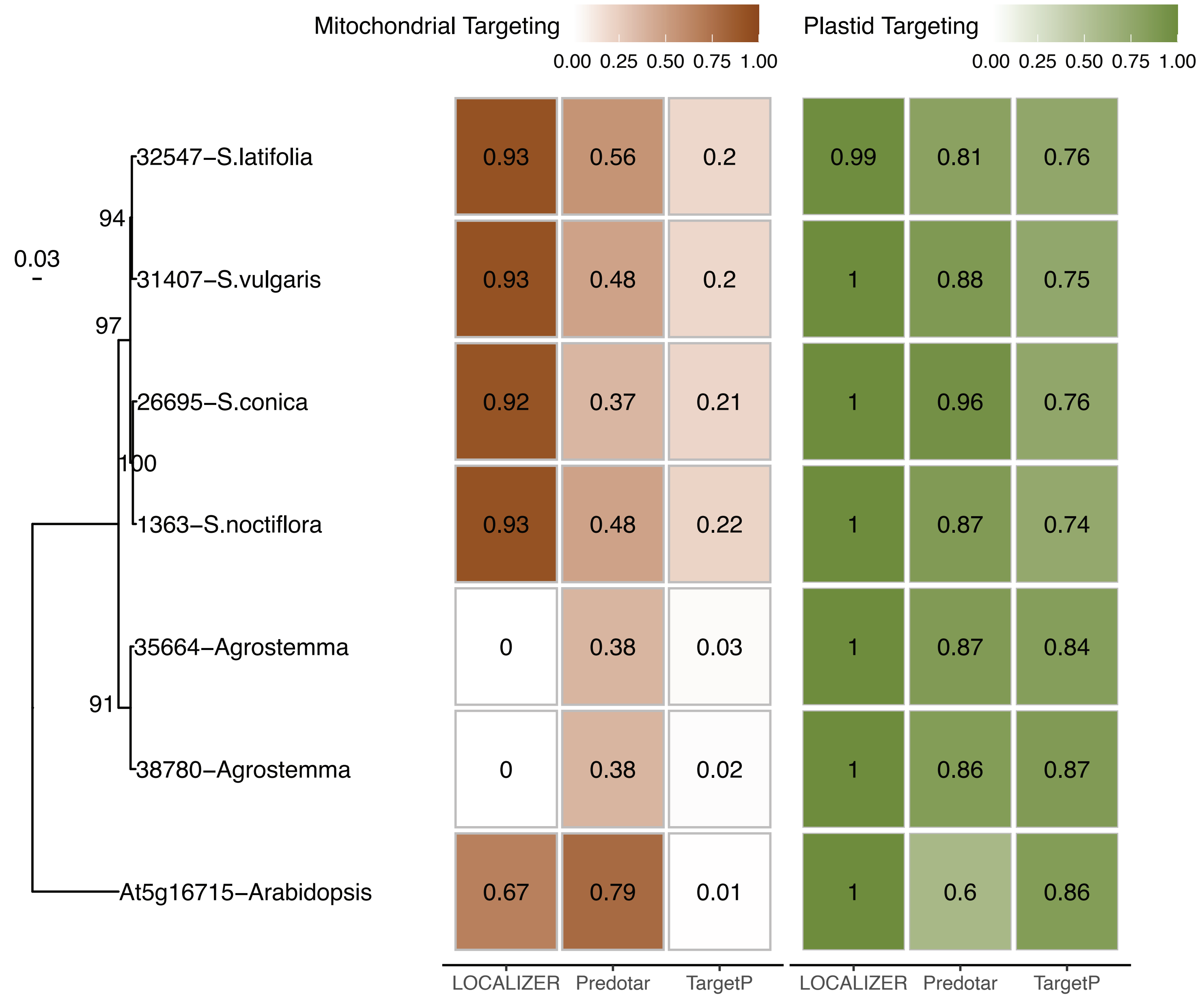

### Supplemental Figure 21

#### Colors for amino acids

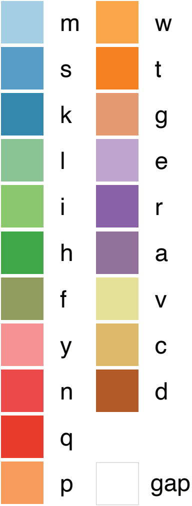
