## Supplemental Table 1 for "Rewiring of aminoacyl-tRNA synthetase localization and interactions in plants with extensive mitochondrial tRNA gene loss"

**Supplementary table 1**| Iso-Seq transcript coverage for identified aaRS genes. Only transcripts with coverage of two or more Iso-Seq reads were retained for analysis (i.e., singleton transcripts were discarded). Single Sileneae transcripts often mapped to multiple A. thaliana aaRS genes because of complex gene histories, including lineage-specific gene duplication events. Organelle localizations marked with * have been experimentally shown (Duchêne et al. 2005). Cytosolic localization were inferred based on the lack of a transit peptide, and a ? indicates that a transit peptide with potential mitochondrial targeting is present, but experimental localization studies were inconclusive (Duchêne et al. 2005) .

| **aaRS** | **Mapped TAIR gene** | **Localization in *A. thaliana*** | **Iso-Seq transcript coverage** | | | | |
|---|---|---|---|---|---|---|---|
| | | | ***Agrostemma*** | ***S. conica*** | ***S. latifolia*** | ***S. noctiflora*** | ***S. vulgaris*** |
| **AlaRS** | At1g50200 | Cytosol/Mitochondria*/Plastid* | 1084 | 786 | 2630 | 60 | 1617 |
| **AlaRS** | At5g22800 | Mitochondria*/Plastid* | 189 | 228 | 143 | 9 | 365 |
| **ArgRS** | At1g66530:At4g26300 | Cytosol:Plastid* | 190 | 139 | 265 | 54 | 511 |
| **AsnRS** | At1g70980:At5g56680 | Cytosol:Cytosol | 135 | 85 | 206 | 42 | 208 |
| **AsnRS** | At4g17300 | Mitochondria*/Plastid* | 68 | 48 | 67 | 14 | 40 |
| **AsnRS** | At3g07420 | Cytosol | 23 | 0 | 0 | 4 | 2 |
| **AspRS** | At4g26870:At4g31180 | Cytosol:Cytosol | 165 | 62 | 114 | 82 | 140 |
| **AspRS** | At4g33760 | Mitochondria*/Plastid* | 44 | 37 | 37 | 6 | 61 |
| **CysRS** | At3g56300:At5g38830:At2g31170 | Cytosol:Cytosol:Mitochondria*/Plastid* | 135 | 102 | 229 | 93 | 116 |
| **GlnRS** | At1g25350 | Cytosol | 640 | 194 | 301 | 65 | 554 |
| **GluRS** | At5g26710 | Cytosol | 139 | 510 | 342 | 97 | 268 |
| **GluRS** | At5g64050 | Mitochondria*/Plastid* | 92 | 48 | 130 | 35 | 86 |
| **GlyRS** | At1g29870:At1g29880:At3g44740 | Cytosol:Mitochondria*:Cytosol | 896 | 445 | 1225 | 90 | 552 |
| **GlyRS** | At3g48110 | Mitochondria*/Plastid* | 114 | 73 | 165 | 6 | 225 |
| **HisRS** | At3g02760 | Cytosol | 525 | 221 | 357 | 11 | 468 |
| **HisRS** | At3g46100 | Mitochondria*/Plastid* | 62 | 23 | 36 | 19 | 29 |
| **IleRS** | At4g10320 | Cytosol | 713 | 687 | 812 | 31 | 684 |
| **IleRS** | At5g49030 | ?/Plastid* | 472 | 258 | 498 | 6 | 331 |
| **LeuRS** | At1g09620 | Cytosol/Mitochondria* | 1441 | 732 | 829 | 27 | 1672 |
| **LeuRS** | At4g04350 | Plastid* | 90 | 27 | 77 | 2 | 94 |
| **LysRS** | At3g13490 | Mitochondria*/Plastid* | 107 | 148 | 199 | 21 | 376 |
| **LysRS** | At3g11710 | Cytosol | 172 | 102 | 501 | 61 | 314 |
| **MetRS** | At3g55400 | Mitochondria*/Plastid* | 197 | 11 | 77 | 10 | 51 |
| **MetRS** | At2g40660 | Cytosol | 53 | 13 | 19 | 33 | 26 |
| **MetRS** | At4g13780 | Cytosol | 202 | 258 | 162 | 62 | 363 |
| **PheRS** | At3g58140 | Mitochondria*/Plastid* | 56 | 22 | 33 | 23 | 35 |
| **PheRS** | At1g72550 | Cytosol | 191 | 63 | 140 | 37 | 293 |
| **PheRS** | At4g39280 | Cytosol | 81 | 118 | 161 | 69 | 165 |
| **ProRS** | At3g62120 | Cytosol | 133 | 233 | 384 | 116 | 356 |
| **ProRS** | At5g52520 | Mitochondria*/Plastid* | 48 | 51 | 73 | 45 | 64 |
| **SerRS** | At5g27470 | Cytosol | 167 | 67 | 68 | 98 | 187 |
| **SerRS** | At1g11870 | Mitochondria*/Plastid* | 35 | 3 | 20 | 0 | 17 |
| **ThrRS** | At2g04842 | Mitochondria*/Plastid* | 120 | 155 | 65 | 53 | 135 |
| **ThrRS** | At1g17960:At5g26830 | Cytosol:Mitochondria* | 390 | 238 | 384 | 66 | 184 |
| **TrpRS** | At3g04600 | Cytosol | 29 | 21 | 24 | 32 | 47 |
| **TrpRS** | At2g25840 | Mitochondria*/Plastid* | 19 | 8 | 5 | 6 | 18 |
| **TyrRS** | At1g28350:At2g33840 | Cytosol:Cytosol | 326 | 12 | 97 | 14 | 189 |
