## Supplemental Table 3 for "Rewiring of aminoacyl-tRNA synthetase localization and interactions in plants with extensive mitochondrial tRNA gene loss"

**Supplementary table 2**| Yields from Iso-Seq libraries.

| Species | CCSs | FLNCs | Clusters (HQ) |
|---|---:|---:|---:|
| *A. githago* | 2034058 | 2029458 | 153293 |
| *S. conica* | 1535114 | 1525890 | 94736 |
| *S. latifolia* | 1584689 | 1580565 | 122274 |
| *S. vulgaris* | 1765441 | 1762081 | 126377 |
